# Measures of epitope binding degeneracy from T cell receptor repertoires

**DOI:** 10.1101/2022.07.25.501373

**Authors:** Andreas Mayer, Curtis G. Callan

## Abstract

Adaptive immunity is driven by specific binding of hyper-variable receptors to diverse molecular targets. The sequence diversity of receptors and targets are both individually known but, because multiple receptors can recognize the same target, a measure of the effective ‘functional’ diversity of the human immune system has remained elusive. Here, we show that sequence near-coincidences within T cell receptors that bind specific epitopes provide a new window into this problem, and allow the quantification of how binding probability co-varies with sequence. We find that near-coincidence statistics within epitope-specific repertoires imply a measure of binding degeneracy to amino acid changes in receptor sequence that is consistent across disparate experiments. Paired data on both chains of the heterodimeric receptor are particularly revealing since simultaneous near-coincidences are rare and we show how they can be exploited to estimate the number of epitope responses that created the memory compartment. In addition, we find that paired-chain coincidences are strongly suppressed across donors with different human leukocyte antigens, evidence for a central role of antigen-driven selection in making paired chain receptors public. These results demonstrate the power of coincidence analysis to reveal the sequence determinants of epitope binding in receptor repertoires.

---

Which epitopes are recognized by an individual’s T cells? The specificity of T cells is encoded genetically in the loci coding for the hypervariable loops of the T cell receptor (TCR) chains [1], and thus in principle reading out the immune repertoire by sequencing provides the information to answer this question [2–4]. Yet, deciphering the complex sequence ‘code’ for the many-to-many mapping between TCRs and peptide-major histocompability complexes (pMHCs) remains an open problem [5].

Aspects of this code are coming into view thanks to data from multiple experimental approaches. Structural studies have revealed the spatial arrangements in which TCRs bind pMHCs [6–11]. Mutagenesis experiments [12, 13] have provided early evidence that some amino acid substitutions in TCRs maintain or even increase binding affinity to a given epitope. Such local degeneracy of the binding code has been confirmed more recently by sequencing of epitope-specific groups of TCRs [14–21], and sequence patterns in these datasets are now used in machine learning approaches to predict further binders to the same epitope [22–26].

Direct experiment can, however, examine only a minute fraction of all the possible binding combinations, due to the enormous diversity of potential receptors and epitopes: more than 10^12^ different peptides [27] are presented on 1000s of human MHC alleles [28] to up to 10^61^ possible TCRs [29] generated by the recombination machinery. As a result, rules that generalize across epitopes would be of utmost utility, but TCR diversity has made it difficult to find such rules.

To address this problem, we here introduce a statistical framework that quantifies the sequence degeneracy of receptors that bind to a common target via the enhanced frequencies of sequence coincidences in epitope-specific TCR repertoires as compared to their frequencies in suitably chosen ‘background‘ repertoires. The specific repertoires we study can be created in a controlled way in an experiment, or can arise organically, as when a memory compartment is formed in response to an infection. Generalizing the analysis to inexact coincidences (pairs of sequences with high sequence similarity), we find that they, too, are enhanced in epitope-specific repertoires. We demonstrate mathematically that the ratio of near-coincidence probabilities between data and background, as a function of sequence distance, is a direct measure for how specificity is correlated across sequence space.

Applying this framework to epitope-specific T cell repertoires that have been acquired in different ways [14–17] reveals a common coincidence enhancement signature of specific binding across disparate experiments. We relate this signature to the existence of a typical average local binding degeneracy, defined as the fraction of the available sequence neighbors of a specific T cell receptor (available in the sense of being present in a natural repertoire) that will also bind to the same pMHC. In addition, we see a weaker version of this signature in paired chain repertoires that have not been subjected to explicit *ex vivo* enrichment for epitope-specific T cells [30]. We exploit this observation in two ways: we provide clear evidence that this signature is associated with MHC presentation of antigen by demonstrating that coincidences between different donors are strongly affected by the overlap between their human leukocyte antigen (HLA) types; in addition, after some mathematical analysis, we use it to quantify the effective functional diversity of the memory repertoire, in the sense of an estimated number of epitope recognition events it records. Taken together, these results illustrate how coincidence analysis can help to quantitatively address immunological questions whose answers have so far remained elusive.

## I. OVERVIEW OF ANALYSIS STRATEGY

We illustrate the broad strategy of our approach on a repertoire of CD8^+^ T cells specific to an Epstein-Barr Virus peptide presented on HLA-A*02:01. The data is from Dash et al. [14] and was obtained using single cell receptor sequencing of tetramer-sorted T cells binding the specific pMHC.

Fig. 1A shows a clustering by pairwise amino acid sequence distance of all distinct nucleotide sequence clones. In this visualization, each position in the heatmap records the sequence distance Δ between the amino acid sequences of a pair of distinct T cell clones. TCRs are heterodimeric, and the heatmaps above (below) the diagonal record distances between the *β* (*α*) hypervariable complementary-determining region 3 (CDR3) loops of the sequence pair. Clustering is based on the sum of distances between *α* and *β* chains. Here, and throughout this work, we define sequence distances as the minimal number of edits (insertions, deletion, or substitutions) that change one sequence into another, known as the Levenshtein distance. We only consider sequence distances between CDR3 loop for simplicity, but the mathematical framework we develop is general and could also be used with distance measures that include other hypervariable receptor regions. By clones we mean lineages of cells that go back to the same ancestral recombination event, which we define in practice based on nucleotide sequence identity. A zero distance pair arises when due to convergent recombination two distinct nucleotide sequences have the same amino acid translation. We chose to ignore the number of times a given nucleotide sequence is sampled, as clone sizes also reflect TCR-independent lineage differences [31, 32]. Instead, we analyze convergent selection imposed on distinct clones with the same or similar TCR as a stringent measure of epitope-driven functional selection. In the experiments that we consider in this manuscript, TCRs are selected for binding to a specific pMHC ligand, and our analysis quantifies the imprint of this filtering funnel on TCR sequence statistics. We use the word “selection” to refer to this filtering process, which is distinct from, and not to be confused with, thymic selection.

**Figure 1:**
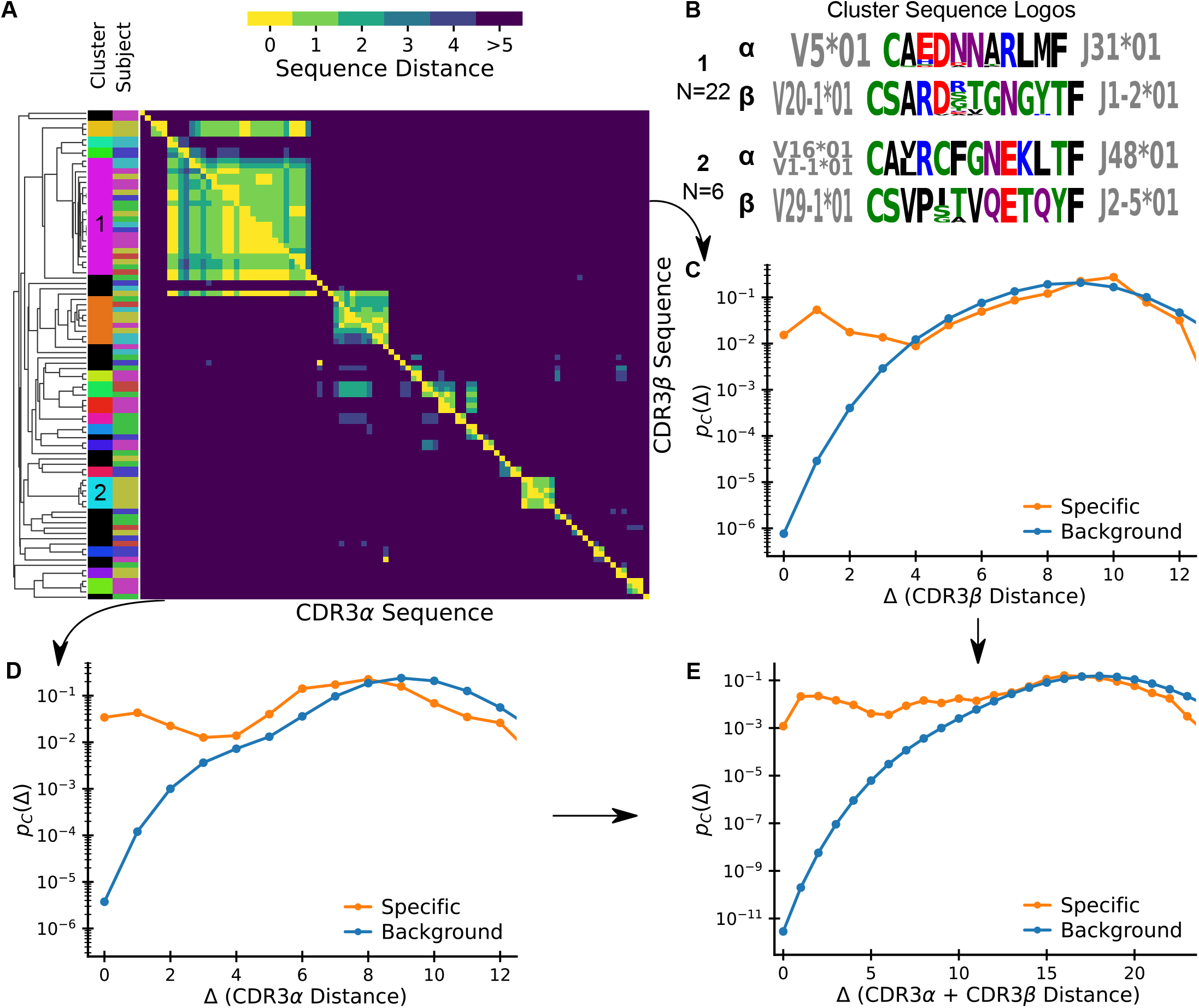
Patterns of sequence similarity within an epitope-specific repertoire. (A) Sequence-similarity clustermap of TCRs binding to an Epstein-Barr Virus epitope as obtained by single cell TCR sequencing following tetramer sorting (Data: Dash et al. [14], antigen BMLF). Lower (upper) triangle shows pairwise distances of CDR3*α* (CDR3*β*) sequences. Sequences are ordered by average linkage hierarchical clustering based on summed *αβ* distance. Columns on the left show subject of origin and cluster assignment; sequences not belonging to a cluster based on a cutoff distance of 6 are shown in black. (B) Sequence logos for two clusters of specific sequences. Amino acids are colored by their chemical properties, and V and J gene usage within the cluster is displayed alongside the logo. (C-E) Normalized histograms of pairwise distances between (C) CDR3*β*, (D) CDR3*α*, and (E) CDR3*αβ* sequences specific to the epitope show vastly increased sequence similarity relative to background expectations.

Fig. 1A allows some direct conclusions about important features of TCR-pMHC binding code: First, it highlights the remarkable sequence similarity among specific TCRs and it shows that this similarity also holds for TCRs from different donors. Second, it shows that there are several clusters of sequences differing by a few substitutions from each other, plus a substantial number of isolated sequences that differ from all other sequences by many substitutions. Fig. 1B shows sequence logos for two prominent clusters. Interestingly, they are quite different from each other, even when accounting for chemical similarity of amino acids. This suggests clusters might represent broad structurally distinct binding solutions, each with local residue degeneracy. This view is supported by the V and J gene usage, which is highly restricted within each cluster but non-overlapping between them. Third, it demonstrates that chain-pairing is biased even among specific binders as similarity on one chain is often associated with similarity on the other chain.

To compare statistics of sequence similarity across epitope targets, we next compress the off-diagonal elements of the clustermap into a normalized pairwise distance histogram that we denote by *p_C_*(Δ). We normalize coincidences by *N* (*N* – 1)/2, the number of possible pairs (i.e. upper diagonal elements in the matrix), so that *p_C_*(Δ) is a probability distribution on Δ. Figs. 1C,D show the histograms for *α* and *β* chains, respectively. Fig 1E shows the histogram for the complete *αβ*-TCR, with paired chain sequence distance defined as the sum of distances of both chains. These normalized pairwise distance distributions are the basic element of our analysis framework. We also plot the *p_C_*(Δ) distributions derived from bulk sequencing of a “background” sample as a proxy for the expected distribution prior to selection. We use sequencing data from Minervina et al. [16] of total peripheral blood mononuclear cells (PBMCs) from a healthy individual for these background curves for *α* and *β* chains. For the paired chain background curve we currently lack sufficiently deeply sequenced datasets. Fortunately, previous studies have concluded that *α* and *β* chain gene usages are largely uncorrelated [30, 33, 34], so we use the convolution of the *α* chain and *β* chain distributions from Minervina et al. [16] as a plausible paired chain background prior to selection. In section VII we will present further evidence supporting the use of this assumption.

The central observation is that *p_C_*(Δ) is orders of magnitude larger in epitope-specific repertoires than the corresponding background for small Δ. Exact coincidence frequencies are in excess by surprisingly large factors (∼ 10^9^ and ∼ 10^4^ for paired and unpaired chains, respectively). This excess extends to near-coincidences, but interestingly, for large enough Δ, the selected and the background values of *p_C_*(Δ) approach each other. The manner in which their ratio falls to unity will turn out to be roughly the same across different types of experiments, an intriguing fact that points to shared underlying biophysical rules of specific binding.

## II. THEORY OF COINCIDENCE ANALYSIS

### A. Definitions and statistical estimation

The T cell clones that enter the immune repertoire are drawn from a background distribution *P*(*σ*) over all possible nucleotide sequences *σ* that code for the TCR hypervariable chains. This distribution summarizes the statistics of the recombination process by which the receptor coding genes are rearranged, and it is known that probabilities of individual sequences range over many orders of magnitude [35]. Experimentally, clones are identified by distinct nucleotide sequences, and coincidences (exact or near) are defined by the corresponding amino acid sequence (since that is what determines functional identity or similarity). Generation probabilities are such that it is unlikely that two separate T cell generation events will give the same nucleotide sequence, but it is less uncommon for them to give the same CDR3 amino acid sequence. Therefore the practical limitation of identifying clones by distinct nucleotide sequences instead of recombination events introduces only minimal bias. The normalized histogram of pairwise distances defined operationally in the previous section is then an empirical estimate of coincidence probabilities, more formally defined as

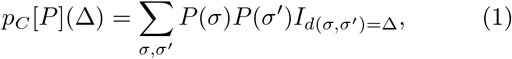

where *I* is the indicator function, the sum is over independent pairs of nucleotide sequences, and *d*(*σ, σ*′) is the sequence distance between the amino acid translations of the sequences.

Given the diversity of TCRs it is surprising that we are able to find any coincidences in small epitope-specific repertoires. The occurrence of coincidences at sample sizes much smaller than the space of all sequences is connected to the “birthday problem” in probability theory [36, 37]: In a sample of *N* distinct sequences there are *N* (*N* – 1)/2 distinct pairs, and the expected number of pairs at distance Δ is thus *p_C_*(Δ)*N*(*N* – 1)/2. This means that we can estimate normalized pair probabilities *p_C_*(Δ) ∼ 10^−3^ using repertoires of only *N* ∼ 10^2^ sequences. This is fortunate since it is precisely this combination of orders of magnitude that we encounter when we estimate *p_C_*(Δ) from epitope-specific repertoires at small values of Δ (Fig. 1C-E).

### B. Intuition for why coincidences increase in epitope-specific repertoires

To gain intuition, we define a probability distribution on amino acid sequences by marginalizing over nucleotide sequences, 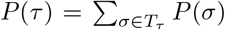, where *T_τ_* is the set of sequences that translate to amino acid sequence *τ*. In this notation, we can give an alternative definition of the exact coincidence probability (the value of Eqn. 1 at Δ = 0) as

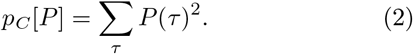

This expression is Simpson’s diversity index from ecology [36]. Its inverse 1*/p_C_* is known as a true diversity, an estimate of an effective number of species present in a population. Here, amino acid receptor sequences take the role of species, which means *p_C_* is an index of the diversity of amino acid sequences coded for by the different clones in the repertoire. Only some receptors bind an epitope, thus we expect epitope-specific repertoires to have lower diversity. This provides an intuitive explanation why *p_C_*, the inverse of true diversity, increases with selection. From this perspective, Eqn. 1 represents a generalization of Simpson’s index to a similarity-weighted measure of diversity [38]. As epitope-specific repertoires consist of TCRs with similar sequences, we expect similarity-weighted diversity to also be restricted. This in turn helps rationalize why *p_C_*(Δ) is increased in epitope-specific repertoires for some range of small Δ. A central point of this paper is that a great deal of information is contained in the generalization of Simpson’s index to inexact coincidences.

To develop this intuition further, let us represent T cells with distinct nucleotide sequences as nodes in a graph, and connect pairs of clones with the same TCR amino acid sequence with a link. Fig. 2A displays such a graph representation for 100 notional background T cells, together with the result of selecting half of them according to two different protocols. The probability that a randomly chosen pair of nodes are linked is equal to *p_C_* = 2|*E*|/(|*V*|(|*V*| – 1)), where |*E*| is the number of edges and |*V*| is the number of vertices. The pre-selection repertoire is shown in the left panel, where links were arbitrarily chosen such that *p_C_* = 0.02. The middle and right panels show the results of two selection protocols mimicking random subsampling and epitope-specific sorting, respectively: selecting nodes with probability 1/2, ignoring linkage (center), or selecting clusters of nodes with probability 1/2 (right). When selecting cells at random, the coincidence probability *p_C_* = 0.02 is unchanged: the mean number of linked pairs decreases by a factor 4, but so does the total number of possible node pairs. Selecting clusters in contrast, implies that the number of edges decreases by only a factor 2. Normalizing by the total number of node pairs, the coincidence probability increases two-fold to *p_C_* = 0.04. The selection of connected clusters mimics sorting by epitope-specificity, in the sense that cells belonging to the same clonotype, defined by identical amino acid sequence, all share the same specificity.

**Figure 2:**
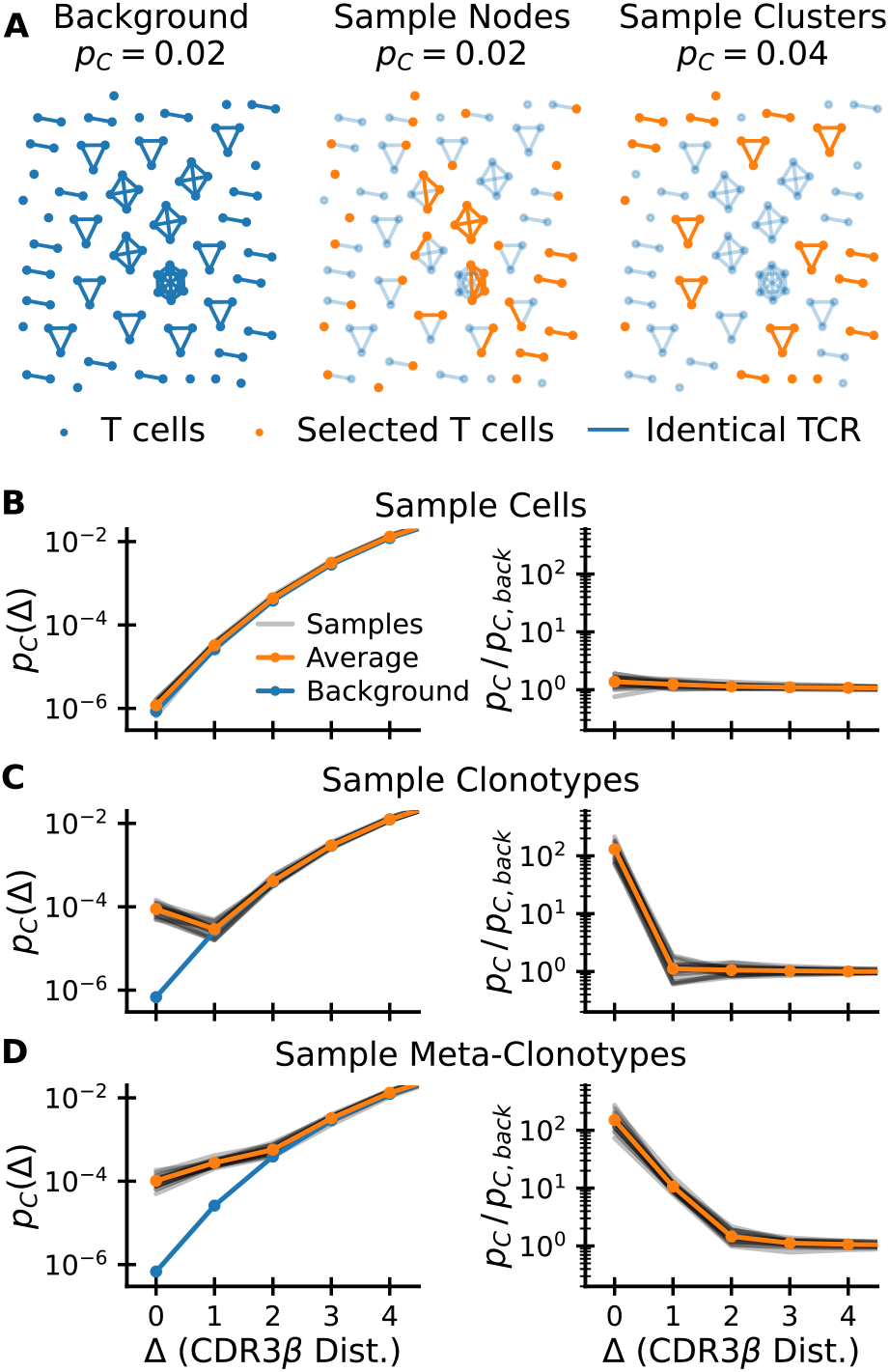
How selection increases coincidences. (A) How different selection procedures change the graph of sequence neighbors. Cells (nodes) in a background graph (left) are connected by edges if they share an identical TCR. Random sampling of nodes (middle) does not change the coincidence probability. Random sampling of clusters (right) increases the coincidence probability. Selected nodes and links in orange; unselected background nodes in light blue. (B-D) Coincidence probabilities for synthetic data generated by selecting 1% of cells (B), 1% of amino acid clonotypes (C), and 1% of meta-clonotypes (generated by including 10% of neighbors of each selected sequence, see text) (D) at random. These random selection protocols act on a a background CDR3*β* repertoire (data from Minervina et al. [16]). The grey lines show estimates for 20 repetitions of the sampling procedure, and the orange line their average.

### C. Formal analysis

We now mathematically derive how coincidence probabilities change when specific TCRs are identified within a larger pool. We analyze this as follows: let *Q*(*σ*), normalized by ⟨*Q*(*σ*)⟩_*P*(*σ*)_ = 1, be a selection factor that characterizes whether sequence *σ* meets the chosen selection condition. The distribution of selected sequences is then *Q*(*σ*)*P* (*σ*). As we derive in Appendix A the coincidence distributions of the two ensembles are related via the cross-moments of the selection factors,

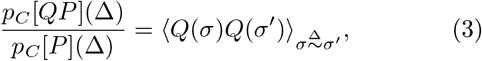

where 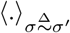 indicates that the average is calculated over random pairs of sequences at distance Δ, i.e. over the distribution *P* (*σ, σ*′ *d*(*σ, σ*′) = Δ).

To gain intuition we consider a simple class of selection functions of relevance to antigen-specific selection, where *Q* weights equally a specific subset of sequences and gives zero weight to all others:

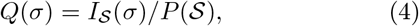

where *P* (*S*) = ∑_*σ*∈*S*_ *P*(*σ*) is the fraction of all clones (i.e. distinct nucleotide sequences) that are specific to the epitope in question. Given the statistical process that created the background repertoire, any given background sequence has an expected number of ‘neighbors’ at sequence distance Δ; if the sequence in question is selected, we can ask what fraction *f_σ_*(Δ) of its neighbors at distance Δ are also selected. Plugging Eqn. 4 into Eqn. 3 we find that the coincidence enhancement ratio is proportional to the average of that fraction over the selected sequences 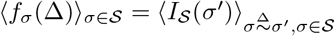:

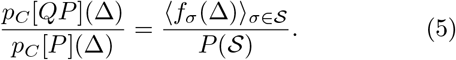

Note that ⟨*f_σ_*(Δ = 0) _*σ*∈*S*_ = 1 because specific binding only depends on amino acid sequence, so that all exact coincidences with a selected sequence must also be selected. Thus, the increase in exact coincidence probability is inversely proportional to the selection fraction *P*(*S*). If the selection fraction is small, the coincidence ratio is large, in line with the interpretation of this ratio as a measure of the strength of selection. What follows is a direct way to estimate the average number of specific neighbors:

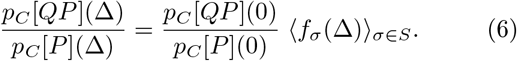

How coincidence ratios decrease with distance Δ is thus a measure of the average sequence degeneracy of specific binding. Applying this equation to experimental data will allow us to estimate this fundamental quantity. In comparing with data the empirical coincidence distribution within an epitope-specific repertoire is our measure of *p_C_*[*QP*], and *p_C_*[*P*] is determined from a background set of sequences. To simplify notations we will thus refer to their ratio as *p_C_*/*p_C,back_*.

### D. Simulation of selection on real data

To make the preceding formal analysis concrete we next turned to numerical simulation of selection of sequences from a realistic background T cell repertoire. To get intuition of the effect of selection by a generic pMHC complex at a gross statistical level, we filter sequences from a background dataset of approximately 10^5^ CDR3*β* sequences taken from whole blood (data from [16]) according to different random sampling protocols.

We first compare selecting random cells (Fig. 2B) with selecting random clonotypes (Fig. 2C), in each instance selecting 1% of sequences. For the former, apart from statistical noise, *p_C_*(Δ) is the same for the selected set as for the background. For the latter, the exact coincidence frequency increases hundredfold. This increase corresponds to the inverse of the selection fraction *P*(*S*) ∼ 10^−2^, exactly as predicted by Eqn. 3. Such random selection of clonotypes was used successfully in Elhanati et al. [39] to predict TCR sharing numbers among a large number of human individuals. However, for Δ ≠ 0 coincidence frequencies do not differ from the background (in contrast to empirical data, such as Fig. 2C).

We thus next sought to incorporate sequence correlation in selection between similar amino acid sequences to model the local degeneracy in antigen recognition apparent in Fig. 1A. To this end, for each selected sequence *σ*, we also select a fraction *p_corr_* of sequences that are within sequence distance Δ_*corr*_ from *σ*. The construction of such a sequence-correlated random selection model is somewhat subtle as a naive scheme oversamples sequences with many neighbors. We derived a corrected sampling scheme explained in Appendix D that overcomes this bias. The results of such a selection of metaclonotypes for Δ_*corr*_ = 1 and *p_corr_* = 0.1 are shown in Fig. 2D. As expected, sequence correlations lead to an enhancement of *p_C_*(Δ) over background that extends to near coincidences. Also, the selection enhancement ratio changes by a factor of ∼ 0.1 (the value of *p_corr_*) between Δ = 0 and Δ = 1, in accord with our expectation from Eqn. 6.

We note from these illustrations that the the enhancement ratio *p_C_*(Δ)*/p_C,back_*(Δ) (plotted in the right-hand columns of Fig. 2B-D) gives a particularly direct diagnostic of the nature and strength of the selection that acts on the background. We will use it in the next sections to put a wide range of experimental data into a common framework.

## III. COMMON FEATURES OF SELECTION ACROSS MULTIPLE DATASETS

We now use the lens of coincidence analysis to examine a broad set of experimental datasets that use different assays to select T cell repertoires specific to epitopes from different sources (details in Methods and Data) [14–17, 30]. Our analysis of these diverse datasets (Fig. 3) reveals striking similarities in the functional dependence of excess coincidences on sequence distance, together with wide variation in the magnitude of the enhancement of coincidence frequencies over background.

**Figure 3:**
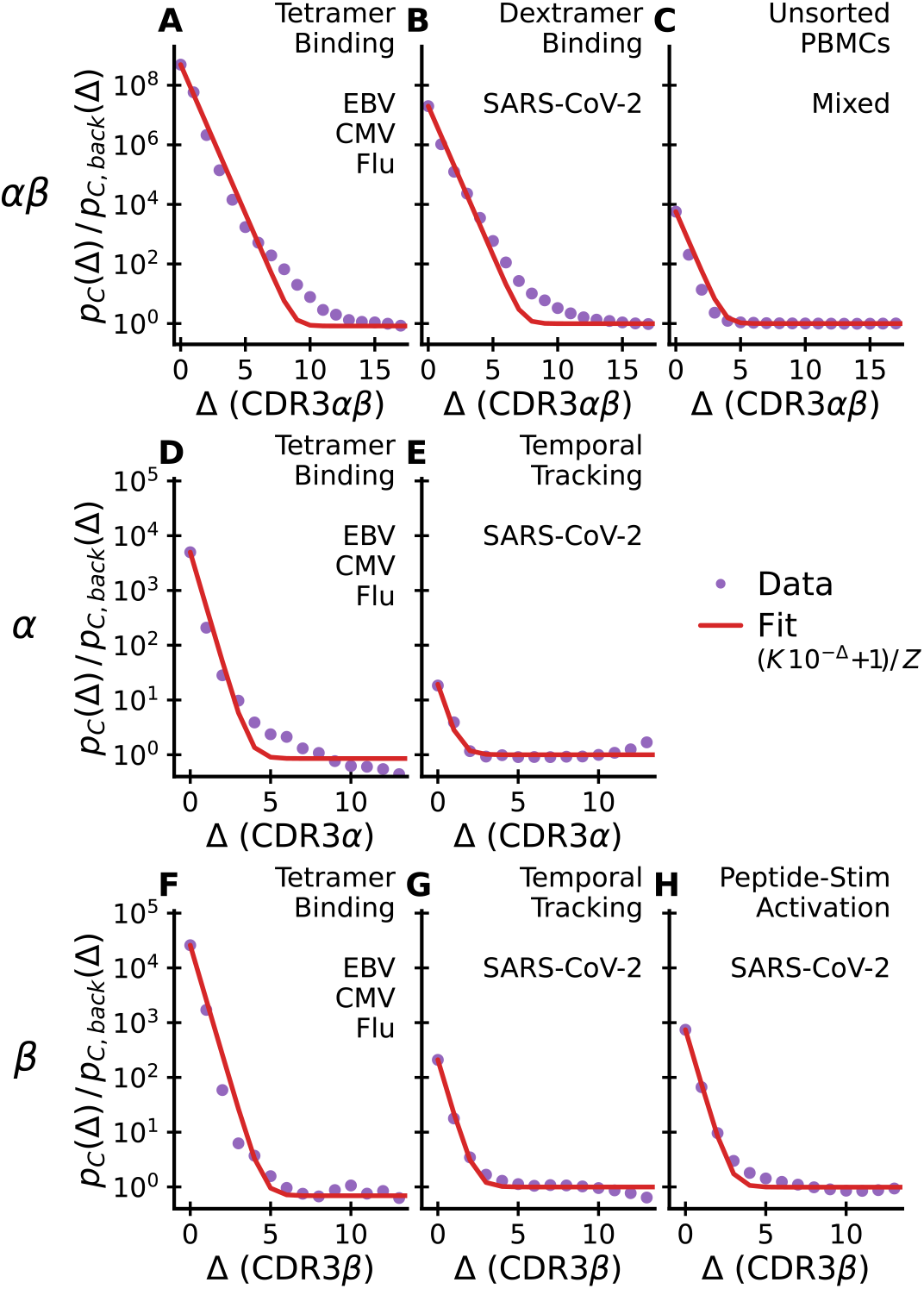
Excess coincidences follow a common functional form across experiments. Sequence similarity of specific T cells for paired *αβ*-chain repertoires (top row), *α*-chain repertoires (middle row) and *β*-chain repertoires (lower row) compared with background expectations. In each panel the assay type used to enrich for epitope-specific T cells and the antigen source are noted in the upper right. Panel C is special as analyzed TCRs are from unsorted blood and have not been explicitly selected for binding to a specific epitope. A common reference curve is plotted for visual guidance. Its parameter K is set equal to the empirical value at Δ = 0. *Z* is determined by normalization. Datasets: A,D,F - Dash et al. [14]; E,G - Minervina et al. [16]; H - Nolan et al. [15]; B - Minervina et al. [17]; C - Tanno et al. [30].

We first apply coincidence analysis to paired chain data from Dash et al. [14] (Fig. 3A), Minervina et al. [17] (Fig. 3B), and Tanno et al. [30] (Fig. 3C), taking the distance between two paired sequences to be the sum of distances between the two chains. Minervina et al. sequenced paired-chain *αβ* TCRs that were determined by DNA-barcoded MHC dextramers to have specificity to chosen SARS-CoV-2 epitopes, while Tanno et al. provides a large dataset of paired-chain total T cell repertoires that have not been directly subjected to *ex vivo* selection. We compute the coincidence probability ratio *p_C_*(Δ)*/p_C,back_*(Δ) against a synthetic background computationally constructed from single chain data under an independent pairing assumption, as described previously.

We next apply coincidence analysis to the single chain data from Nolan et al. [15] (Fig. 3H) and Minervina et al. [16] (Fig. 3E,G). Nolan et al. sequenced *β*-chain sequences of T cells selected (by passage through the MIRA pipeline [40]) for recognizing individual peptides in a broad panel of peptides from the SARS-Cov-2 genome, while Minervina et al. identified *α* and *β* chain sequences of T cells that responded dynamically during the SARS-Cov-2 infection of two human subjects using longitudinal sequencing. As a comparison we also analyze single chain sequences from the Dash et al. [14] paired chain dataset across the three studied viral epitopes, ignoring the chain pairing (Fig. 3D,F). For each repertoire, we compute the coincidence probability ratio *p_C_*(Δ)*/p_C,back_*(Δ) against background bulk sequences of the same chain. To smooth out variability, we then average over epitopes or subjects, respectively.

Together, these analyses highlight major differences across chains and experiments (Fig. 3, rows and columns, respectively) in how much coincidence probabilities are increased relative to background, *p_C_*(Δ)*/p_C,back_*(Δ) at small Δ. The fold increase for sequence identity (Δ = 0) is highest in paired chain tetramer-sorted repertoires against immunodominant epitopes of common viruses (Fig. 3A), and decreases from this value when chains are considered separately (Fig. 3 2nd and 3rd row) or in sequence repertoires identified by other assays (Fig. 3 2nd and 3rd column). We will provide a potential mechanistic explanation for some of these differences in Section VI.

There are also some striking common features to note. First, the analyses show that, for small Δ and across experiments, the excess coincidence ratio declines from its value at Δ = 0 at a similar exponential rate; second, across all datasets, coincidence rates reduce to those of the background for distances substantially less than the mean distance between sequences in the background. In other words, the statistical differences between selected repertoires and the background are limited to small sequence distances Δ. The red curves plot a simple parametric function (specified in the legend) that captures the two key features: it interpolates between an initial exponential decrease by roughly one power of ten per unit increase in Δ and asymptotes to a constant. The parameter *K* is set to the value of excess coincidences at Δ = 0, and the parameter *Z* is determined self-consistently by normalization. Without any additional fitting parameters the reference curve is in good agreement with the empirical data across all experiments, highlighting their similarity.

The exponential falloff for small Δ is a quantitative measure of binding degeneracy with respect to small sequence changes. According to Eqn. 6 the observed common falloff rate means that, on average, about one tenth of the Δ = 1 sequence neighbors of a T cell that recognizes an epitope will also recognize the same epitope (and roughly one percent of the Δ = 2 neighbors, etc). This degree of sequence degeneracy is observed both for *α*-chains (Fig. 3D,E) and *β*-chains (Fig. 3F-H). Note that this analysis relates to the fraction of available sequence neighbors, i.e. those present in the pool before sorting for specificity in accord with the TCR generation probabilities and sample size, and takes into account only the CDR3 region and not other hypervariable regions. The observation that this parameter agrees across experiments and chains is striking, and suggests it is a fundamental biophysical feature of TCR-pMHC binding interactions.

## IV. DIVERSITY OF BOTH CHAINS AND THEIR PAIRING IS RESTRICTED IN SPECIFIC TCRS

Epitope-specific repertoires sequenced at the paired-chain level can be used to quantify the relative contribution to binding specificity of the two chains. Figs. 3D,F show that there is, on average, a strong diversity restriction (as measured by excess coincidences) for both chains individually due to epitope selection. If the selected chains could be freely paired without affecting specificity, then the overall excess coincidence factor for paired chains would be the product of the factors for the individual chains (as discussed in Appendix C). In fact, Fig. 3 A shows that paired chain coincidences are more frequent than this expectation by perhaps as much as a factor 10 (out of an overall increase by a factor of ∼ 10^9^). For further insight, we repeated the analysis separately for each individual epitope (Fig. 4): the paired chain selection factor is in each instance the product of two large factors due to selection of the *β* and *α* chains individually times a smaller factor that arises from restricting pairing among the selected sets of chains, and there is only limited variation in the contributions of the three terms across epitopes. These results show why paired chain information is essential for accurately predicting the specificity of a TCR. An important correlate of the strong restriction of diversity within epitope-specific repertoires is that when fixing one chain the other shows only very limited variation: As shown in Fig. 1 paired chain coincidences are nearly as frequent as coincidences on either chain alone. A related phenomenon was recently described comparing naive and memory antibodies [41], and termed chain coherence. Our analyses suggest that such coherence also occurs for TCRs.

**Figure 4:**
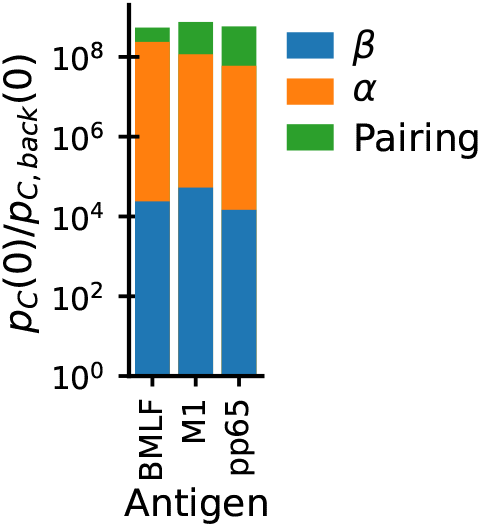
Epitope binding restricts diversity of both chains individually, and also restricts their pairing. Bar chart shows the decomposition of paired chain exact coincidence probability ratios (Fig. 3A) for individual epitopes in the dataset from Dash et al. [14] into contributions from selection of *α* chains (Fig. 3D) and *β* (Fig. 3F) individually (blue,orange), plus a smaller contribution from restricting the pairing of the two chains (green).

## V. THE SELECTION SIGNATURE CONSTRAINS THE BINDING LANDSCAPE

What are minimal features of a T cell-epitope binding landscape that can explain the coincidence enhancement signature? To explore this question we go beyond the random selection models considered in Fig. 2 and treat selection more realistically as due to sequence-dependent binding. This exercise could be carried out at many levels of sophistication [42, 43], but we will focus on a simple, schematic, and analytically tractable model for TCR-pMHC interactions. In what follows, we sketch the model and the conclusions we draw from it. Details are presented SI Sec. F.

We model TCRs as random amino acid strings of fixed length k=6 (corresponding to the mean number of hyper-variable residues within a typical CDR3 loop). Background TCR sequences are generated by drawing 6 amino acids independently at random from the *q* = 20 amino acids. The set of TCRs binding to a specific pMHC is specified by a sequence logo, or motif, condition: at each of the k variable positions, we require that the residue lie in a randomly chosen subset of size *c* ≤ *q* of the amino acids (a different subset at each position).

Calculating the coincidence enhancement factor for a particular epitope and binding motif reduces to a combinatorial excercise in this model, with the result:

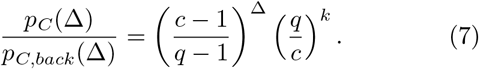

This expression reproduces the exponential falloff of excess near-coincidences with Δ that is seen in real data. The falloff rate depends on the number of allowed amino acids *c* at each position, with *c* ∼ 3 amino acids per position reproducing the empirical rate.

However, this expression does not capture the second observation in the empirical data, namely, that beyond a certain sequence distance Δ, the enhancement ratio asymptotes to a roughly constant value. To address this, we recall that Fig. 1 strongly suggests that there are multiple “solutions” to the problem of recognizing a given epitope. Sequence similarity between TCRs binding in different m anners i s e xpected t o b e l ow, t hus t he existence of multiple solutions might explain the flattening of the coincidence probabilities for large Δ. We thus extend our binding model to incorporate this idea: For each epitope, let there be *M* different randomly chosen motifs and declare that a T cell recognizes the epitope if any of the motifs are satisfied. T cells selected by this model are a mixture of those selected by the individual motifs. Applying results for coincidences in mixture distributions (derived in Appendix B), we obtain an analytical prediction for excess coincidences:

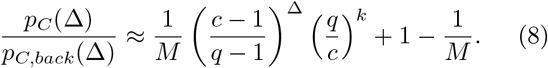

Fig. 5 displays this analytical result for different values of M. In addition it shows the almost identical results of numerical simulations of the model with a more realistic non-uniform amino acid usage (drawn according to the amino acid usage in CDR3*α* hypervariable chains reported in [16]). The key observation is that, for multiple motifs, the ratio *p_C_/p_C,back_* both shows exponential decay for small Δ and asymptotes to a constant (close to unity) as Δ approaches the maximum possible value in this setup, Δ = 6.

**Figure 5:**
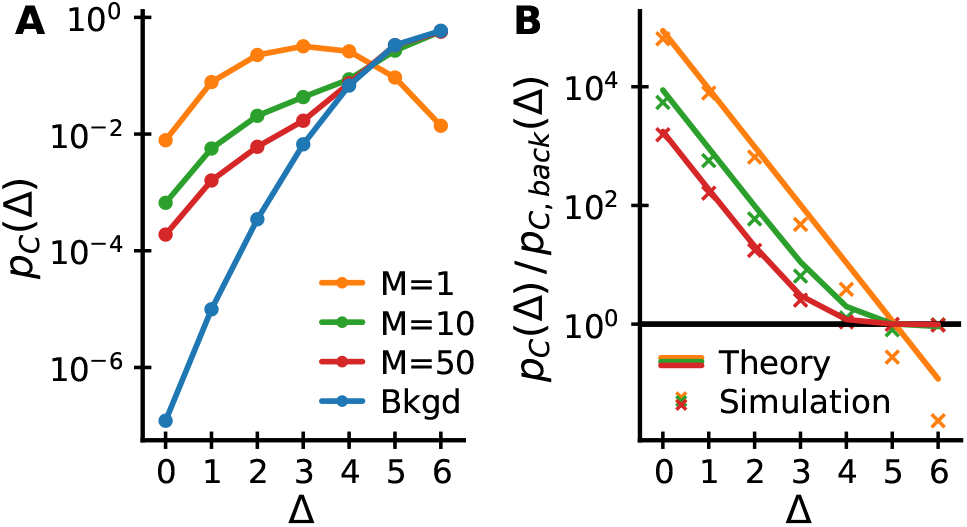
Coincidences in a mixture of motifs model. (A) Coincidence probabilities and (B) coincidence probability ratios to background for simulated data generated from a mixture of motifs model with different numbers of motifs *M* and *c* = 3. (B) also shows analytical expectations from Eqn. 8 (lines), which agree well with the numerical results (crosses). The model reproduces key features of the empirical data: *p_C_/p_C,back_* decays exponentially for small Δ and asymptotes to a constant for large Δ at sufficiently large *M*.

## VI. FUNCTIONAL DIVERSITY LINKS COINCIDENCES ACROSS SCALES

We now revisit the intriguing observation of a selection-like signature in paired chain sequencing data from whole blood (specifically, the coincidence enhancement displayed in Fig. 3C). In Fig. 6 we compare coincidence frequencies obtained from direct paired chain sequencing of blood samples with coincidence frequencies among multimer-sorted T cells that recognize individual epitopes. We note that coincidences within multimer-sorted repertoires exceed those in blood samples by four orders of magnitude. Also, the comparison with sorted memory and naive repertoires shows that coincidences in the total repertoire are primarily driven by memory cells. Bearing in mind that the whole blood coincidence analysis compares sequences within and between all the memory sub-compartments created by past infections, we hypothesize that the coincidences in whole blood reflect high-levels of sequence similarity among groups of memory cells selected in response to specific epitopes encountered in the past. Intuitively, we then expect coincidences in whole blood to depend on the diversity of the memory repertoire, i.e. on how many different epitope exposures the immune system is remembering. To make this intuition quantitative we develop a mathematical formalism to predict coincidences in mixture distributions.

**Figure 6:**
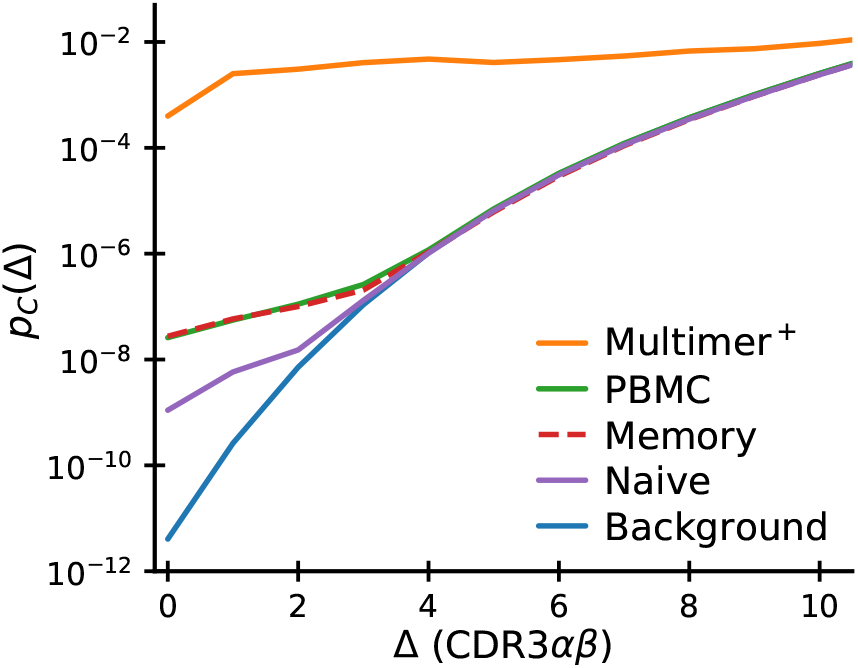
Comparison of near-coincidence probabilities across paired-chain datasets. The highest values come from TCR repertoires specific to individual epitopes (solid orange curve: average over epitopes studied in Dash et al. [14] and Minervina et al. [17]). Paired-chain sequencing of whole blood (green), sorted CD4^+^ memory (dashed red) and CD4^+^ naive (purple) repertoires (data averaged over subjects from Tanno et al. [30]) give much smaller values. Background coincidence probabilities (calculated assuming independent chain pairing) are shown in blue. See text for discussion of the large difference in coincidence probabilities between repertoires.

We propose to model TCRs in an individual’s memory compartment as a mixture distribution over the set Π of peptide-MHC complexes (pMHCs) that have driven past immune responses in that individual. For each *π* ∈ Π, there is a distribution of T cell sequences *P*(*σ*|*π*) that target *π*. The distribution of TCRs in the memory compartment will then be the mixture distribution

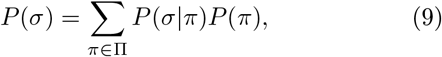

where *P*(*π*) is the proportion of all TCRs selected for binding to pMHC *π*. The coincidence probability for mixtures can be calculated using the following mixture decomposition theorem, which we derive in Appendix B:

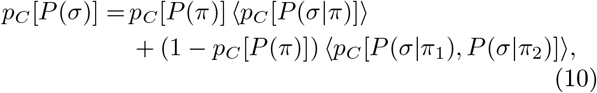

where the averages are over *P*(*π*|*π*_1_ = *π*_2_ = *π*) and *P*(*π*_1_, *π*_2_ *π*_1_ ≠ *π*_2_), respectively. It is noteworthy that such an exact decomposition of coincidence probabilities in mixtures exists. For example, no equivalent formula exists for Shannon entropy, an alternative measure of diversity, which has led to long-running debates within ecology about the decomposition of diversity in pooled communities [44–46].

Eqn. 10 is a sum of two non-negative terms, each of which can be given an intuitive interpretation. We recall that the probability of exact coincidence is the probability with which two randomly chosen sequences *σ*_1_ and *σ*_2_ are coding for the same TCR, *p_C_*[*P*(*σ*)] = 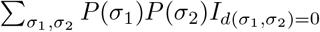. The decomposition formula then represents a conditioning on the mixture identity for *σ*_1_ and *σ*_2_: The overall probability of coincidence is a weighted mean of average within-group coincidence probabilities (first term) and of average between-group coincidence probabilities (second term). The relative weight given to within group comparisons is given by the probability with which two randomly chosen elements come from the same group, i.e. the coincidence probability of the group assignments *p_C_*[*P*(*π*)] (defined in the sense of Eqn. 2).

Multimer sorting followed by sequencing gives draws from *P*(*σ*|*π*) for specific pMHCs *π* [14, 17], and this data can be used to estimate the average within-epitope-group coincidence probability ⟨*p_C_*[*P*(*σ*|*π*)]⟩. Absent better information, we shall assume that the average value *p_C_*[*P*(*σ*|*π*)] ∼ 10^−4^ in these experiments is the typical order of magnitude for all epitopes. We further assume that the between-epitope-group term in Eqn. 10 is negligible. Then the only remaining quantity is *p_C_* [*P*(*π*)], the Simpson diversity of the set of epitope-specific groups within the repertoire. Putting the numbers together we obtain an effective diversity 1*/p_C_* [*P*(*π*)] ∼ 10^4^, a not implausible value for the pMHC diversity of a memory compartment.

In other words, the large ratio between coincidence frequencies in a repertoire selected *ex vivo* by an individual pMHC complex and the coincidence frequencies in the memory compartment as a whole is informative about the number of epitope recognition events that created the memory compartment in the first place. While the precise numbers are likely to change as more comprehensive data becomes available, the calculation above gives a clear recipe to settle the question of how functionally diverse our immune repertoire is. More broadly, mixture averaging also likely explains why coincidence probabilities among longitudinally identified TCRs (presumably specific to multiple immunodominant epitopes) a re lower than among TCRs specific to individual epitopes (Fig. 3E vs. 3D and Fig. 3G vs. 3F,H).

## VII. HLA OVERLAP DETERMINES COINCIDENCES BETWEEN DONORS

How many TCRs are shared between donors? In previous studies of T cell repertoires, there has been much interest in such shared sequences, on the grounds that such ‘public’ sequences may point towards common pathogen exposures [39, 47]. Since in order to mount a common response to a pathogen epitope, two subjects must not only share (up to near-coincidence) T cells with the same TCR, but must also share an MHC molecule on which the epitope can be presented, we expect more T cell sharing between donors that share HLA alleles. In line with this expectation, Tanno et al. [30] observed an association between exact sharing of paired *αβ* TCRs and the number of shared HLA alleles. By our logic, it makes sense to broaden the definition of public T cells to those that are nearly coincident across donors, and present at rates well above an appropriately estimated background. We will thus revisit the analysis by Tanno and co-workers by applying our coincidence analysis framework to their dataset. Specifically, we c alculate the h istogram of sequence distances between TCRs drawn from pairs of repertoires, and ask how the strength of any selection signal depends on the similarity of HLA type between the two repertoires.

We grouped subject pairs by HLA overlap defined as *J* = |*A* ∩ *B*|/ max(|*A*|, |*B*|), where *A* and *B* are the sets of HLA alleles in the two subjects. The overlap ranges between *J* = 1 for identical twins, to *J* = 0 if there is no common HLA allele. We also applied additional filtering steps to control for confounding factors (Appendix E). To mimic the filtering applied to intra-sample analyses of the data from Tanno et al. [30], we did not count coincidences where either chain had exact nucleotide identity. This filtering also allowed us to exclude exact nucleotide coincidences when comparing repertoires of twins. Exact nucleotide-level sharing of full *αβ* TCRs between twins can represent long-lived clones shared via the blood supply during fetal development [48, 49], and is thus not necessarily evidence of convergent selection on the TCRs. Additionally, we removed sequences whose *α*-chain V and J genes match those of two non-canonical T cell subsets, mucosal associated invariant T cells (MAITs) and invariant natural killer T cells (iNKTs), that recognize non-peptide ligands not presented on classical MHC [50].

The results of the analysis are shown in Fig. 7: Near-coincidence probabilities between whole blood repertoires decrease systematically with decreasing HLA overlap (Fig. 7A), and the same trend holds in sorted CD4^+^ memory (Fig. 7B) and CD4^+^ naive cells (Fig. 7C). These HLA-dependent effects are large: exact coincidence probabilities range over two orders of magnitude as HLA overlap varies. This contrasts with prior studies that have found only a small influence of HLA type in single chain repertoires [51]. The interpretation suggested by our earlier analysis (Fig. 4) is that HLA binding requires specific pairs of *α* and *β* chains. To confirm that our observed large effect sizes are compatible with weak signals in single chain repertoires, we constructed synthetic distributions for randomized *αβ* pairings by convolving the single chain distance distributions within HLA overlap groups. The results are shown as dashed lines in Fig. 7 (using the same color coding for the HLA overlap groups as for the real data). They reveal that single chain coincidences are almost independent of HLA overlap, even though this procedure retains the correlations between individual chains and HLA type.

**Figure 7:**
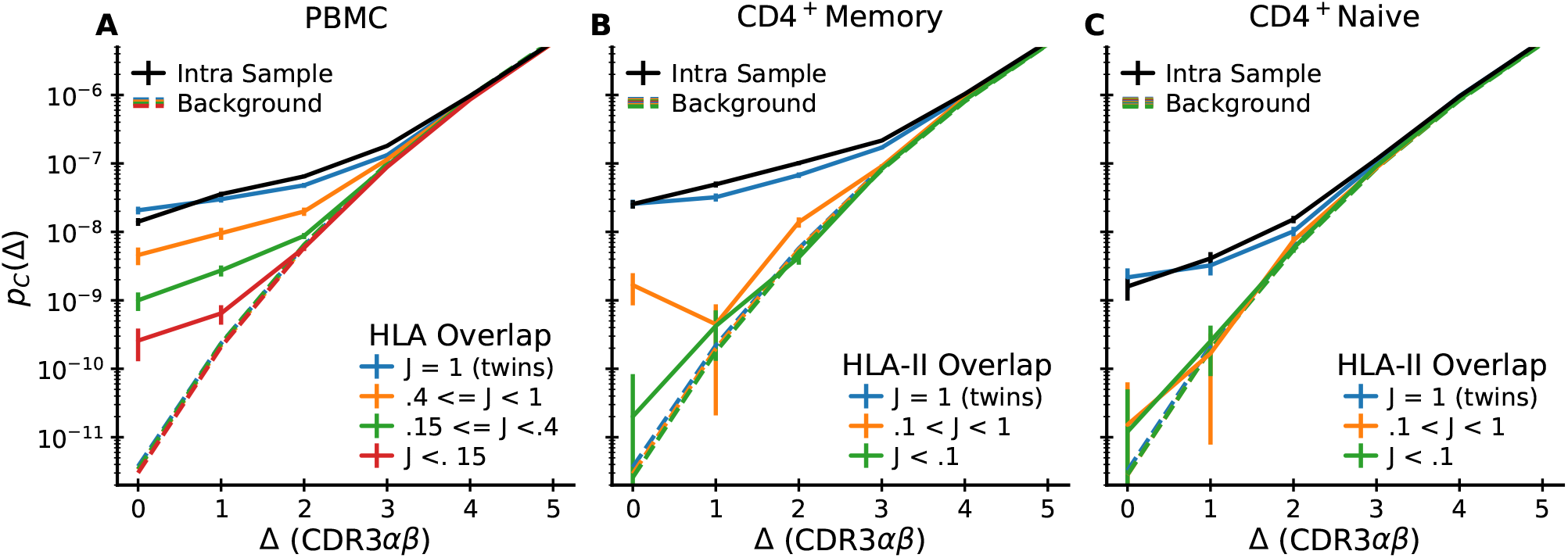
Inter-subject coincidences depend on HLA overlap. Pairwise inter-dataset coincidence frequency analysis for the 15 paired-seq datasets from Tanno et al. [30] grouped by pairwise HLA overlap. A: pairs of unsorted PBMC repertoires; B: pairs of CD4^+^ memory repertoires; C: pairs of CD4^+^ naive repertoires. Each plot shows means over pairs whose HLA overlap lies within the indicated ranges together with estimated standard errors assuming Poisson sampling. For comparison, the mean intra-dataset coincidence distribution is shown in black. Background distributions constructed by scrambling the *α* and *β* chain associations within individuals are shown as dashed curves (colored according to the same HLA overlap code). These curves show no near-coincidence enhancement signal and very weak dependence on HLA overlap class.

The comparison of coincidence probabilities between these different ways of filtering and segregating the data is informative about how different mechanisms might contribute to chain pairing biases. First, Fig. 7C shows no significant deviations from pairing independence (dashed lines in the figure) across naive cells from non-twin donors. This limits the strength of chain pairing correlations that might arise through pMHC-independent processes, such as VDJ recombination, or from steric and biophysical constraints between chains for protein folding [33, 34]. We note that this finding validates the use of the independent chain pairing assumption for generating background distributions representative of repertoire statistics before selection has acted. Second, Fig. 7C also shows a clear signal of correlated chain pairing in naive cells both intra-sample (black line) and across twin pairs (blue line). This strongly suggests that thymic positive and negative selection substantially contribute to the pairing biases. Third, Fig. 7B shows that within the memory repertoire coincidences between twins occur at remarkably similar rates to the intra-sample coincidence rates, which suggests that memory selection is driven by prevalent pMHCs encountered by both donors (herpesviruses are one potential source of such pMHCs [52]). Alternatively, sequences binding a certain HLA might generally show substantially restricted pairing independently of which peptide is presented [53, 54] – something we will soon be able to test as more epitope-specific repertoires for different peptides presented on the same MHC are characterized. In summary, HLA-dependent selection leads to major biases in the pairing of TCR *α* and *β* chains at the repertoire level, the outcome of a combination of thymic and peripheral selection pressures. As dataset sizes continue to increase, the strategy we have described here provides a strategy for untangling these pressures in detail.

## VIII. DISCUSSION

In this work we have introduced a versatile statistical framework for measuring selection in T cell receptor repertoires. Simply put, we have evidence of selection if the number of exactly (and nearly exactly) coincident receptor sequence pairs in a repertoire is substantially larger than the number that one would find in a reference repertoire. Importantly, we showed that this intuitive notion can be developed into a mathematical theory relating the number of excess coincidences to quantities of direct immunological interest, such as the extent of sequence degeneracy of T cell binding to particular epitopes, or the functional diversity of an individual’s memory repertoire.

We take a probabilistic approach to selection, where each target epitope defines a probability distribution on the unselected, or naive, T cells that make up the immune repertoire. Experiments that query blood samples for binding to a specific pMHC represent a draw from this probability distribution, and experiments that capture T cell responses to multiple targets sample a mixture of distributions over targets. Certain global quantities of immunological interest are averages over these distributions and, in our approach, the experimental data serve to give empirical estimates of these averages. We highlight two salient examples:

First, we quantify the fraction of sequence neighbors of a typical specific sequence that share the same specificity. Our analysis predicts that when varying single amino acids in the hypervariable regions in accord with the TCR generative statistics, roughly one out of ten such changes lead to a receptor that still binds the same target. Across disparate datasets this measure of local recognition degeneracy shows remarkable consistency. We envisage that it can be used to guide bioinformatic clustering methods for finding groups of T cells with common specificity [14, 19, 55], for instance to put data-driven constraints on threshold choices. Importantly, the predicted level of local degeneracy is in rough accord with measured distributions of binding affinity changes between point-mutated TCRs [56, 57], and results from systematic mutational scans of specific binding upon changes in TCR hypervariable regions [58]. To quantitatively compare our results with such scans, it will be necessary to develop a framework for appropriately weighting the exhaustive mutational scanning data by the probability with which mutated TCRs occur in natural repertoires. With the rapid increase in the number of assayed epitopes, another area for future work will be to characterize in detail variation around the average selection strength and binding degeneracy, including for example between TCRs binding MHC-I or MHC-II (most data analyzed in the current study relates to MHC-I binding).

Second, we provided a recipe to quantify the functional diversity of a T cell compartment, as measured by the number of different epitopes that have selected the T cells comprising the compartment. From paired-chain sequencing data on human blood samples [30] we derived a rough estimate of the functional diversity of a typical memory compartment. This coarse-grained functional diversity is orders of magnitude smaller than TCR sequence diversity, which is consistent with the relatively small number of immunodominant epitopes typically targeted in response to individual pathogen infections [59] and theoretical predictions that adaptive immunity learns sparse features of the epitope distribution [60]. Additionally, cumulative coincidence probabilities at different sequence distances should provide a useful measure of repertoire diversity weighted by sequence similarity, a subject of recent interest in the field [38, 57, 61].

Beyond the quantification of functional diversity our analysis of deeply sequenced paired chain repertoires across individuals suggests additional research directions. We identify a substantial number of TCR specificity groups that the data suggest are in large part driven by common epitopes across individuals. Guided by such TCR groups it would be interesting to generalize the recently proposed reverse epitope discovery approach [62, 63] to the repertoire scale: cross-referencing coincident TCRs with other data, such as TCR-epitope databases [20, 64] and computationally predicted HLA binding of putative peptides, might guide the identification of the targets of these groups of T cells. More broadly, as dataset sizes increase an analysis of the dependence of cross-donor coincidence probabilities on which HLAs the two donors share could allow an unbiased apportionment of the immune repertoire selected by different HLA types.

In summary, our results reveal both complexity and predictability in the immune receptor code. The emerging picture is captured schematically in the mixture of motifs model that we have introduced: Epitope-specific repertoires are characterized by globally diverse binding solutions that sometimes share surprisingly little sequence similarity, but also display remarkably consistent signatures of local degeneracy. This picture, if further confirmed in structural studies [8, 9], can help focus future machine learning efforts in this area. The consistent signal of local degeneracy suggests that a promising direction will be to use machine learning to refine metrics, such as TCRdist [14, 55], that can group TCRs specific to a common target within large mixtures. Our framework should be of use in such efforts, as it can readily turn any definition of TCR similarity, not just the simple edit distance we have considered here, into probabilities of shared specificity. The existence of multiple binding solutions, on the other hand, might explain why purely sequence-based models for computationally predicting binding partners of epitopes (i.e. in the absence of any experimentally determined binders) have had limited success [23], and why structural modeling might be needed to resolve the complex sequence determinants of the different binding solutions [65].

## Methods and Data

In this paper, we analyze datasets that represent significantly different approaches, both conceptual and experimental, to creating functionally selected T cell repertoires. They are succinctly described as follows:

The Dash dataset [14] is based on tetramer sorting of CD8^+^ T cells from blood, using three well-studied standard viral epitopes (HLA-A*02:01-BMLF1_280_ (BMLF), HLA-B*07:02-pp65_495_ (pp65), and HLA-A*02:01-M1_58_ (M1)), followed by single-cell TCR sequencing to obtain paired TCR*α* and TCR*β* reads of the captured cells. This protocol was repeated for 32 donors, resulting in a list of 415 paired *αβ* TCRs associated with the three epitopes.

The Minervina 2022 dataset [17] uses DNA-barcoded MHC dextramers to identify T cells specific to 19 SARS-CoV-2 epitopes by sequencing. T cells were identified across a cohort of donors with a varied history of SARS-CoV-2 exposure and vaccination. We focused our analysis on the 8 epitopes for which there are at least 150 characterized *αβ* TCRs each.

The Nolan dataset [15] is obtained by sorting about 3 × 10^7^ T cells from a subject blood sample, then incubating the sorted cells with a cocktail of several hundred SARS-CoV-2 epitopes (chosen for their broad MHC presentability) to uniformly expand clones that recognize any of these epitopes. In a next step, aliquots of the expansion product are incubated with individual epitopes from the cocktail, followed by TCR*β* sequencing to identify T cells that have expanded in this second step in response to individual epitopes. This yields a list of TCR*β* clonotypes that recognize the epitope. This protocol is repeated for blood samples from about a hundred subjects, about a third of whom have had no known exposure to SARS-CoV-2 (“healthy” subjects). Summing over subject samples for each epitope, we get a list of a few tens to a few thousand clonotypes that recognize a given epitope. All told, the dataset is a list of some 10^5^ TCR*β* recombination events that respond to individual SARS-CoV-2 epitopes. We note that the *α* chains associated with each *β* chain are not known, and also that a given epitope may be presented on different MHC molecules in different individuals. To have adequate statistical power, we consider only epitopes from Nolan et al. [15] which are recognized by at least 150 distinct clones and we restrict our analysis to MHC-I epitopes.

The Minervina 2021 dataset [16] is based on a longitudinal study of TCR*β* sequences in the blood of two unrelated subjects who contracted mild COVID-19. Analysis of time-separated samples allowed the identification of T cell clones, whose clone sizes changed significantly in response to infection. We focus on the several hundred CD8^+^ clones, whose size decreased between the peak immune response at 15 days and a post-infection time point at 85 days. The specific epitopes to which these T cells respond are unknown, but they are presumably a subset of the SARS-CoV-2 viral epitopes that provoke the strongest immune response, and therefore constitute an interesting “selected” subset of the T cell repertoire.

The Tanno dataset [30] consists of paired-chain TCRs from a total of fifteen donors, including six pairs of twins. The mean number of reads is about 31,000 (minimum of 7400 and maximum of 69000). For three pairs of twins and three unrelated donors total PBMC samples were sequenced. Sorted CD4^+^ naive (CD45RA^+^, CCR7^+^) and memory (CD45RA^−^) cells were sequenced for three additional twin pairs. All fifteen subjects were HLA typed on the allele level. We used processed data as described in the original study but applied additional filtering steps, the rationale for which is described in Appendix E. For the naive repertoire we also removed any overlap with clonotypes that were also found within the memory repertoire from the same individual. To compare coincidence frequencies across repertoires from different individuals (Fig. 7) we sum the number of coincidences across all comparisons within an HLA overlap bin. We add a pseudocount of 0.1 to the summed counts for visualization purposes, and we display Poisson errorbars as 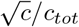 where *c* is the count at a specific distance and *c_tot_* the sum of all counts across distances. These errorbars represent lower bounds, as in addition to counting error there is heterogeneity between individuals.

For all datasets, we filtered out clones whose CDR3 amino acid sequence did not start with the conserved cysteine (C), or end on phenylalanine (F), tryptophan (W), or cysteine (C).

To calculate paired chain background coincidence probability distributions, we randomly associate chains from bulk single chain datasets. For efficient numerical calculation we exploit the fact that such independent pairing leads to coincidence probability distributions for paired chain TCRs that are a convolution of the single chain distributions, 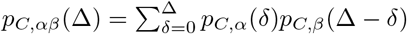.

To generate the sequence logos displayed in Fig. 1 we built on the Python logomaker package [66], adding the ability to also display V and J gene usage. We colored amino acids by their chemical properties using the “chemistry” color scheme.

## Acknowledgements

We are grateful to Yuval Elhanati and Léo Régnier for previous collaboration on the sequence space structure of T cell repertoires, unpublished work that was essential to the development of the ideas underlying the current work. We thank Hidetaka Tanno for providing processed data files and Giulio Isacchini for helpful discussions.

## Author contributions and competing interests

CC and AM conceived research, performed analysis, and wrote the paper. CC and AM declare no competing interests.

## Data and software availability

All the data used in this work has been published and is available online (see the data publications for detailed instructions). We have made an implementation of coincidence analysis available as an open source Python package https://github.com/andim/pyrepseq.

## Significance statement

Adaptive immunity relies on the binding of molecular targets by a few specific T cells out of a highly diverse repertoire. Different T cell receptors can bind the same target, but a quantification of this recognition degeneracy is lacking. Here we develop a statistical approach that links distributions of sequence similarity among T cells of common specificity to how binding probability co-varies with sequence. Applying our method to experimental data we determine the fraction of sequence neighbors of a specific T cell that also bind its target and we estimate how many response groups make up a memory compartment. Our study provides a quantitative framework for identifying the sequence determinants of specific binding, and will thus facilitate the development of repertoire sequencing-based immunodiagnostics.

## Appendix A: Formal treatment of effect of selection on coincidence statistics

## 1. Coincidence changes are related to the cross-moments of the selection factors

We are interested in how the probability of coincidences changes as we modify a base measure *P*(*σ*) with a weighting function *Q*(*σ*) that represents the effect of selection according to

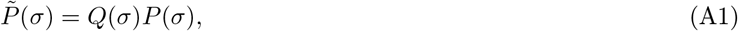

with ⟨*Q*(*σ*)⟩_*P*(*σ*)_ = 1 for normalization of 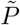. Plugging in 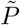 into the definition of the near-coincidence probability, we have

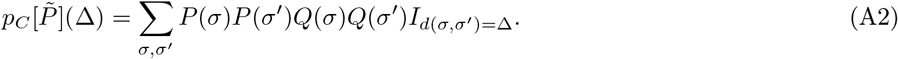

This expression can be rewritten formally as an average over randomly picked pairs of sequences,

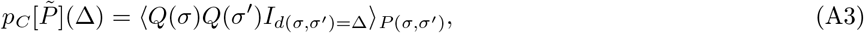

where *P*(*σ, σ*′) = *P*(*σ*)*P*(*σ*′). Only pairs that are at distance Δ contribute to the average, which suggests restricting the average to these pairs. When conditioning on pairs of sequences at distance Δ, the conditional probability of sequences *σ, σ*′ reads

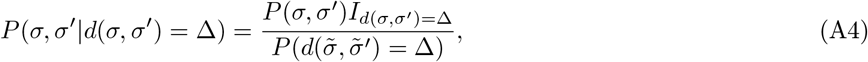

where 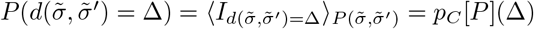 is the near-coincidence probability in the unselected set. The probability of coincidence thus changes upon selection according to

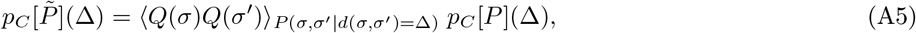

which completes the derivation of Eqn. 3 in the main text.

## 2. Relation to the covariance of selection factors

To gain intuition into Eqn. 3 we can rewrite the first factor on the left hand side in terms of the covariance of selection factors. In general, Cov(*X, Y*) = ⟨*XY* ⟩ − ⟨*X*⟩⟨*Y*⟩. Thus

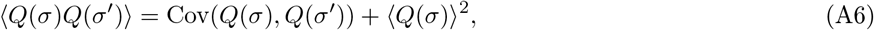

where the covariance is calculated across random pairs at distance Δ, *P*(*σ, σ*′|*d*(*σ, σ*′) = Δ), and the average across the marginal distribution, 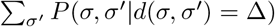. The normalization of 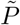 implies ⟨*Q*(*σ*)⟩_*P*(*σ*)_ = 1, hence

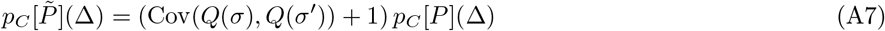

when the average of *Q*(*σ*) over the marginal distribution can be approximated by its simple average over *P*(*σ*). To derive the condition for which Eqn. A7 is exact, we define the local neighbor density around a sequence *σ*,

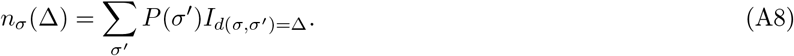

Using this definition we can write

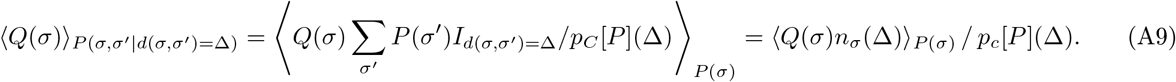

When the probability of selecting a sequence is uncorrelated with its neighbor density, we have

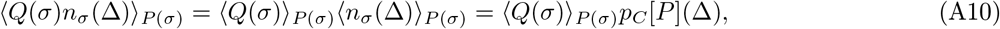

and thus the averages are the same, 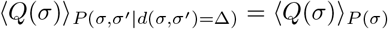 and Eqn. A7 is exact.

## Appendix B: Decomposing coincidences in mixtures

Here we give the formal derivation of Eqn. 10 in the text which describes a situation where the underlying distribution on sequences is a mixture of distributions, each one describing the sequences selected by a particular antigen (or peptide-MHC complex). As discussed in the text, this is a way to describe the memory compartment of the immune system, or the set of TCRs selected by a peptide pool. We define the mixture distribution as

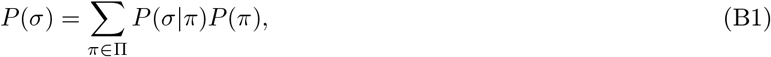

where *P*(*π*) are the mixtures weights with ∑_*π*_ *P*(*π*) = 1. To simplify notations, our derivation will use the exact coincidence definition introduced in Eqn. 2, which uses *P*(*τ*) the marginalized distribution over amino acid sequences *τ*, but we expect the results to hold more generally. We start our derivation by inserting the mixture distribution definition into the formula for the coincidence probability, and split the resulting expression into two terms:

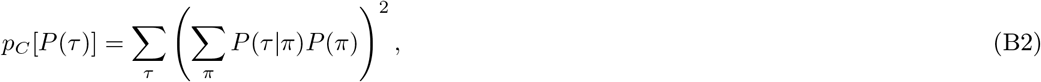

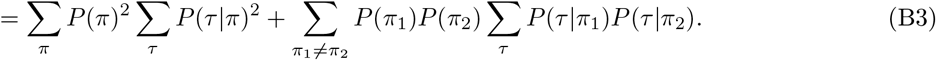

We identify the last factor in the second term as a generalized coincidence probability for samples drawn from two probability distributions,

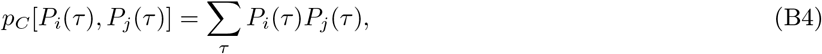

which allows us to rewrite the coincidence probability s

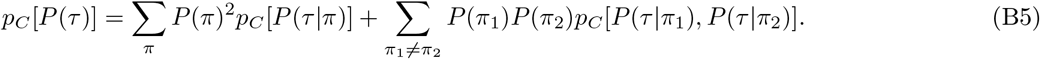

We now further note that

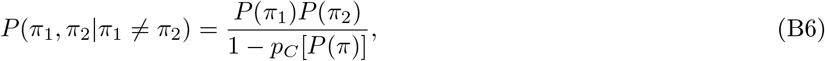

and finally that

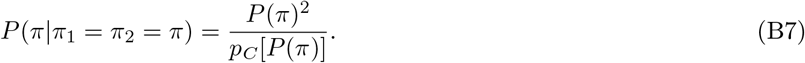

Putting it all together we obtain a rather simple decomposition of the coincidence probability in mixtures:

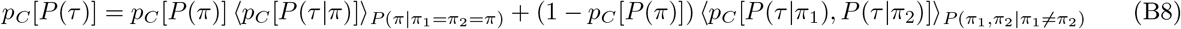

This is equivalent to the equation presented in the main text, expressed there in terms of *P*(*σ*). Let us finally note that we can also generalize Eqn. B4 to inexact coincidences, which defines the generalized cross-sample near-coincidence probabilities,

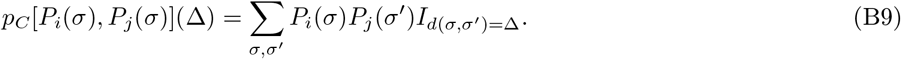

## Appendix C: Decomposing selection on paired chain data

How does selection on the heterodimeric TCR protein restrict diversity on the two constituent chains? To answer this question we start by introducing some notation. The complete clone *σ* is defined by both its *α* chain sequence, which we denote *σ_α_*, and its *β* chain sequence, which we denote *σ_β_*. We define the set of all complete TCRs that bind a specific epitope *S*, such that *σ* ∈ *S* implies the specificity of the receptor encoded by the nucleotide sequence *σ*. The overall selection factor is then equal to

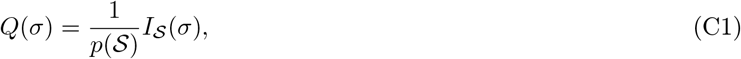

and the exact coincidences in the paired chain data are enriched by a factor *χ_αβ_*,

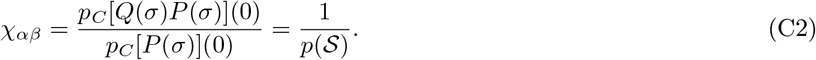

In the following we will ask how this ratio relates to the same ratios calculated for the individuals chains, this is to

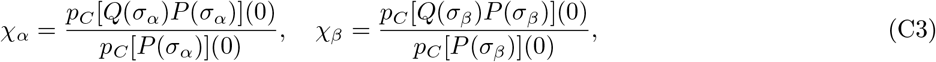

where we will assume independent chain pairing in the background *P*(*σ*) = *P*(*σ_α_*)*P*(*σ_β_*).

Given the full model the selection coefficients for single chains are marginalized averages over all possible choices for the second chain. To calculate these we can rearrange terms in the definition of the marginal single chain post-selection distributions:

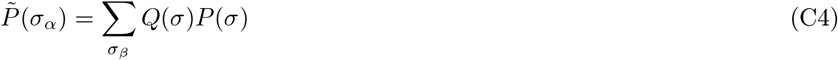

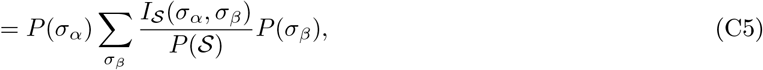

from which we read off

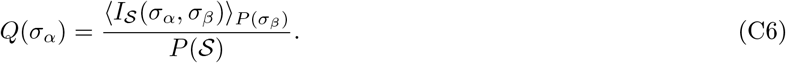

Importantly, we show in the following that *χ_αβ_* = *χ_α_χ_β_*, when there are no chain pairing biases within the epitope-specific set of sequences. To start, note that the set of specific *αβ* sequences is given by the cartesian product

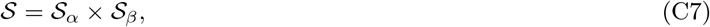

of the single chain sets

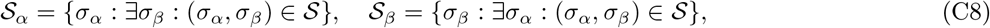

which are composed of all chains that when paired with any of the opposite chains are specific. From these definitions it follows that

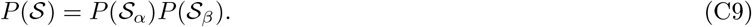

Furthermore some algebra shows

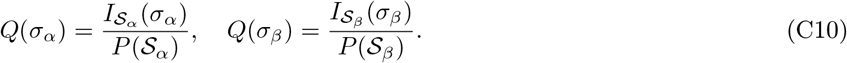

and thus

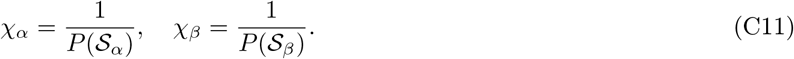

Combining this result with Eqn. C9 we complete the derivation of the equality

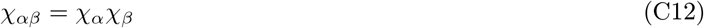

In the general case, the set *S* of all specific sequences is a proper subset of the cartesian product *S*_*α*_ × *S*_*β*_, and thus *P*(*S*) < *P*(*S_α_*)*P*(*S_β_*). We thus expect the coincidence ratio for the paired chain receptors to increase. This motivates using the ratio

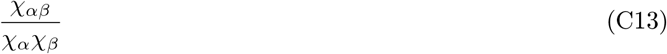

as a measure of pairing biases among the specific receptor chains.

## Appendix D: Random models of sequence-correlated selection

In Sec. II D of the main text we defined a simple random model of sequence-correlated selection on a background repertoire of recombination events (or clones) by a notional epitope as follows: choose at random a fraction (1% in the examples studied) of the CDR3 amino acid sequences that appear in this background and declare that any recombination event with a CDR3 amino acid sequence included in this list is ‘selected’; in addition, let a random fraction (10% in the examples that follow) of all background events with CDR3 amino acid sequence lying at Levenshtein distance one from an element of the selected list be declared to be selected as well. This procedure is motivated by the observation that systematic studies of T cell selection by individual epitopes show a) the existence of multiple unrelated ‘solutions’ in sequence space to the problem of binding a specific epitope, and b) the existence of some degree of sequence degeneracy within individual binding solutions.

This protocol will bias selection toward sequences that have more than the average number of near neighbors in sequence space. While there is nothing wrong in principle with this, systematic studies of selection by multiple epitopes (such as [15]) show little if any such bias. This motivates us to modify the basic random selection procedure to make the probability of selecting a given background sequence independent of the number of its neighbors in sequence space. In what follows, we approximately solve a simplified version of this problem using the language of random graphs, and use that solution to propose a modification of the sequence-correlated random selection algorithm. We then show by concrete example that this modification achieves the desired result in the more demanding context of selection from realistic T cell sequence ensembles.

## 1. Mathematical analysis of the two-step selection algorithm

Consider the following random graph problem: we have a set of *N* nodes, with links between the nodes defined by an adjacency matrix *A_ij_* where *A_ij_* = 1 if node i and j are connected by an edge, and *A_ij_* = 0 otherwise. The nodes represent amino acid sequences in a background repertoire, and links connect sequence distance one sequence neighbors. We define a ‘selected’ subgraph by assigning Ising variables *s_i_* to each node to indicate whether the node is selected (*s_i_* = 1) or not (*s_i_* = 0). Our goal is to find a way of choosing the *s_i_* such that a specified fraction of the nodes have *s_i_* = 1 and the probability that a particular node *i* is selected is independent of the number of links that node participates in. Thus the selected subgraph inherits a subset of the nodes *i* of the original graph and its adjacency matrix is the projection of *A_ij_* on the surviving nodes (no new nodes are created).

We define a two step selection process: We first select a fraction *q*_1_≪1 of the nodes and we then select a fraction *q*_2_ (typically *q*_2_ ≫ *q*_1_) of the links that connect a selected node to one that was not selected; when such a link is selected, we add the originally unselected node to which it connects to the list of selected nodes (i.e. we capture some of the neighbors of a selected node). We take *η_i_* (*θ_ij_*) as independent binary random variables (0 or 1) that describe which nodes (links) are picked during the first (second) step of the selection procedure, respectively and have averages *q*_1_ (*q*_2_) respectively.

The variable *s_i_* that indicates whether a given node was picked in either of the two steps can be written as

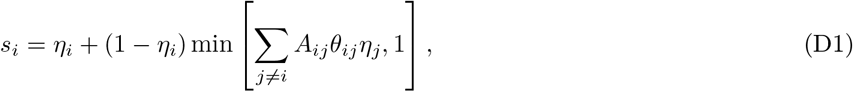

The min function in the second term accounts for the possibility of multiple selection of the unselected node *i* in the link selection step. In practice, this occurs very rarely for the values of the parameters *q*_1,2_ that are of interest to us. We will thus ignore this non-linearity in what follows for tractability. We can calculate various marginals and correlations involving the *s_i_* by averaging over the independent, uncorrelated, binary variables *η_i_* and *θ_ij_*. Doing so, we find

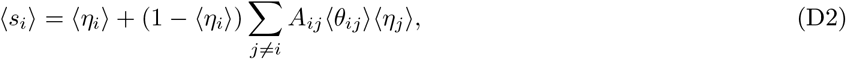

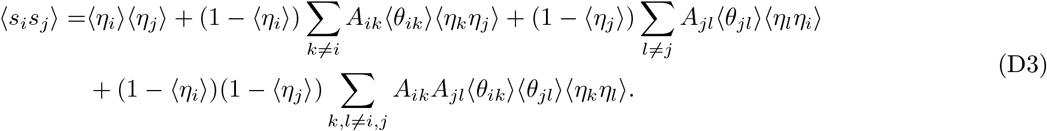

For i ≠ *j*, *η_i_* and *η_j_* are statistically independent by assumption and, because *η_i_* is binary, 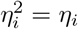. This yields the identity

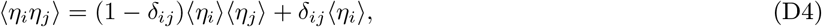

which we can use to simplify the equation for the joint selection probability of a pair of nodes as follows:

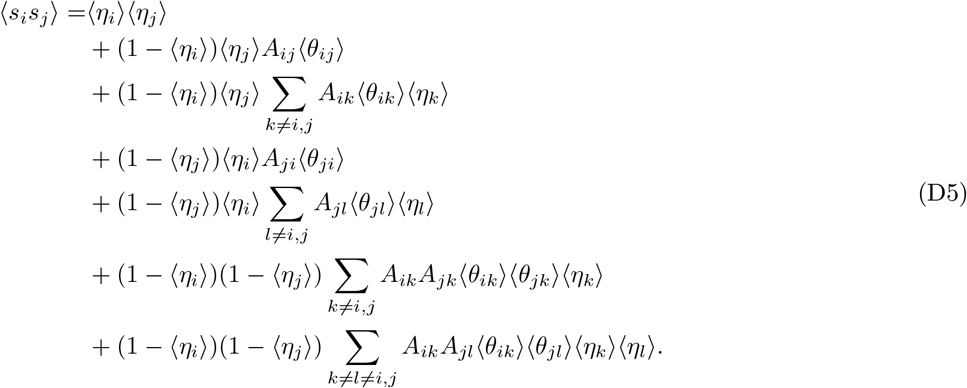

Let us use these expressions to analyze the naive algorithm for sequence-correlated random selection that was described at the beginning of this appendix. In this scheme 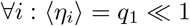 and 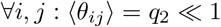. Importantly, for nodes *i*, *j* connected by an edge (so that *A_ij_* = 1), the leading order terms in the above expression for ⟨*s_i_s_j_*⟩ give, for *q*_1_ ≪ *q*_2_ (the case of interest):

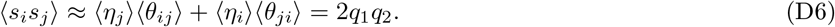

In other words, the procedure allows us to enhance the probability of picking adjacent vertices beyond the independent probability 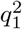 of picking two isolated vertices.

However, as mentioned above, this procedure introduces a bias towards sampling highly connected sequences. More precisely, according to Eqn. D2

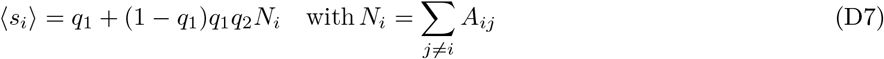

where *N_i_* is the number of edges connecting to the i-th vertex: the more connected nodes *i* will have higher probability of being selected. The idea of the corrected procedure is to make the selection probability in the first step dependent on the neighbor number: ⟨*η_i_*⟩ = *q*_1_(*N_i_*) where *q*_1_(*N*) is a function to be determined. The probability with which node *i* is selected in this procedure can be expressed as

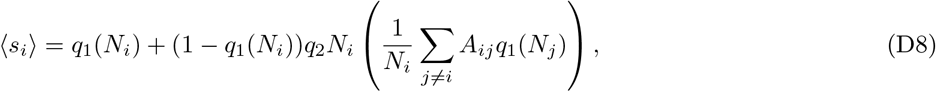

where the term within brackets is the average first step selection probability of all neighboring sequences connected to *i* by a link. To derive an appropriate functional form for *q*_1_(*N_i_*) we make the further assumption that neighboring nodes have the same number of edges on average, so that 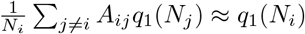. One can then show that the choice of

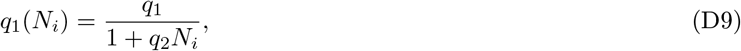

leads to an approximately constant probability of selection, ⟨*s_i_*⟩ ≈ *q*_1_, independent of *i*, if *q*_1_ ≪ 1.

## 2. Numerical tests of the corrected algorithm

How well does proposed neighbor-number corrected selection algorithm work on realistic T cell sequence data? We have applied the algorithm described in the previous paragraphs to background repertoires of approximately 10^5^ sequences of naive CD8^+^ T cells obtained from a human subject (part of the immuneCODE dataset [15]). We generated 100 selected repertoires of ∼ 10^3^ sequences from this background using the neighbor number corrected algorithm. In this background repertoire, the average number of distance-one neighbors of a sequence is close to three and the variance is very large. This analysis is thus a good test for how well the corrected selection algorithm succeeds in equalizing selection probability for nodes with different numbers of neighbors.

In Fig. 8 we plot the probability of selection of a recombination event from the background repertoire, averaged over 100 realizations of the algorithm. We use a fixed selection parameter *q*_1_ = .01 and neighbor selection parameters *q*_2_ ranging from 0.0 to 0.1. We plot the selection fraction as a function of the number of distance one neighbors the selected node has in the background repertoire. The net selection probability except for fluctuations is nearly constant out to neighbor number of ∼ 10, a value that captures 90% of the nodes in the background.

**Figure 8:**
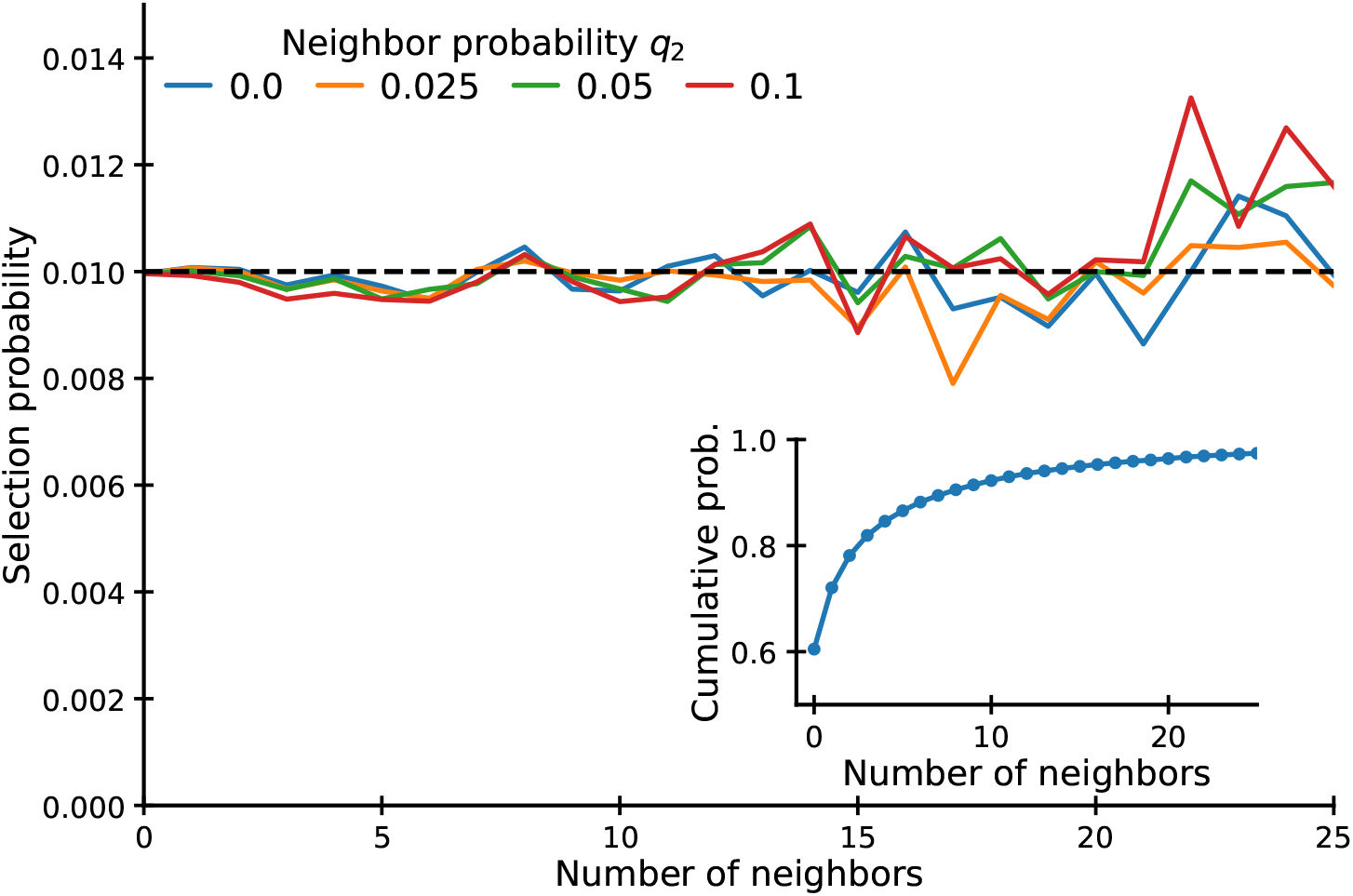
Assessment of the uniformity of random selection from background sequence repertoires across groups of sequences classified by different numbers of Levenshtein distance one neighbors. The plot shows the fraction of each class in the background repertoire that appears in the selected repertoire, averaged over 100 realizations of the selection algorithm.

It is instructive to display the near-coincidence frequency distributions that result from the corrected selection algorithm for different values of *q*_2_ (Fig. 9). As expected from Eqn. 6 the rate of falloff of the coincidence frequency enhancement over background at small sequence distance Δ depends on the parameter *q*_2_ that governs the fraction of the neighbors of a selected sequence that will also be selected.

**Figure 9:**
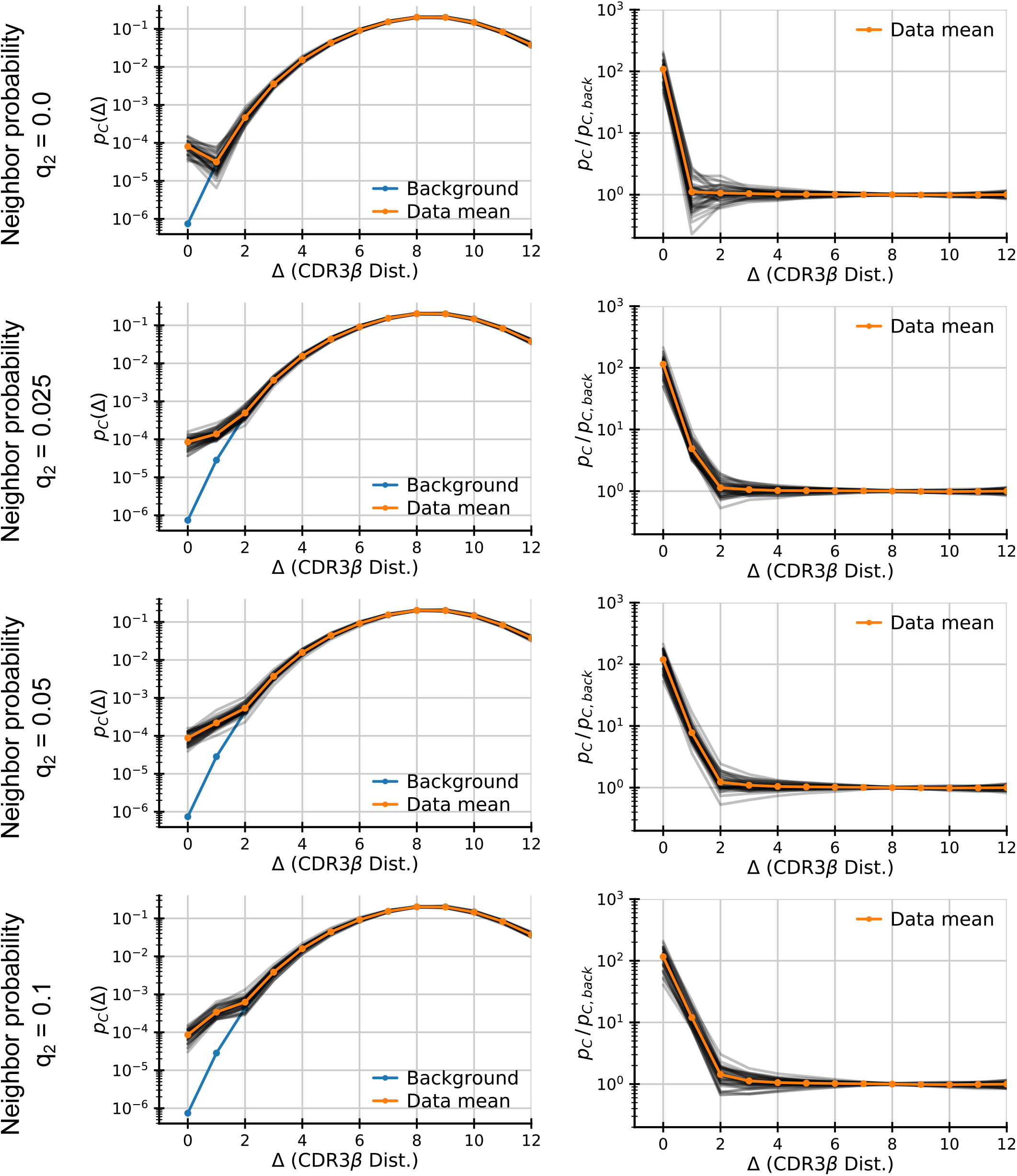
Coincidence frequency distributions for different realizations of the neighbor-corrected random selection algorithm. The black lines give the results of individual random realizations of selection, while the orange curves give averages over the full set of 100 realizations. The key feature to note is the inverse correlation between *q*_2_, the parameter governing the sequence neighbor selection probability, and the initial slope of the log ratio of near-coincidence frequencies to background.

## Appendix E: Analysis of paired chain sequencing data from Tanno et al.

In the following we describe some observations about the nature of sequence coincidences in the paired chain sequencing data from Tanno et al. [30], that have prompted us to perform further filtering steps beyond the analysis pipeline proposed in the previous work.

The experimental protocol uses overlap extension polymerase chain reaction to link both *α* and *β* hypervariable chains before sequencing. Linking predominantly occurs between mRNA from the same single cell as those are captured on the same bead. However, cross-contamination can lead to erroneous pairing of mRNA from different cells. To reduce cross-contamination the authors of [30] clustered CDR3*β* on 95% sequence identity (for typical sequence lengths this is ≤ 2 nts), and kept only the most frequent CDR3*α* sequence from each cluster. No such clustering had been performed for CDR3*α*, and we thus asked to what extent contamination accounts for CDR3*α* sequences associated with multiple CDR3*β* sequences. To assess this we calculated the ratio between CDR3*α* coincidence probabilities on the nucleotide and amino acid level (Fig. 10A). As a comparison we calculated the same ratio for sequences generated using a probabilistic model of VDJ recombination [67] and found that ratios greatly exceed expectations. This finding suggests substantial cross-contamination that might impact downstream analysis. We thus implemented a filtering step clustering CDR3*α* sequences with exact sequencing identity, and kept only the most frequent CDR3*β* sequence from each cluster. Such filtering substantially reduces coincidence probabilities bringing them closer in line with those observed in a bulk-sequenced *α* chain repertoire dataset (Fig. 10B). Variability across subjects is also reduced, suggesting different levels of cross-contamination across samples.

**Figure 10:**
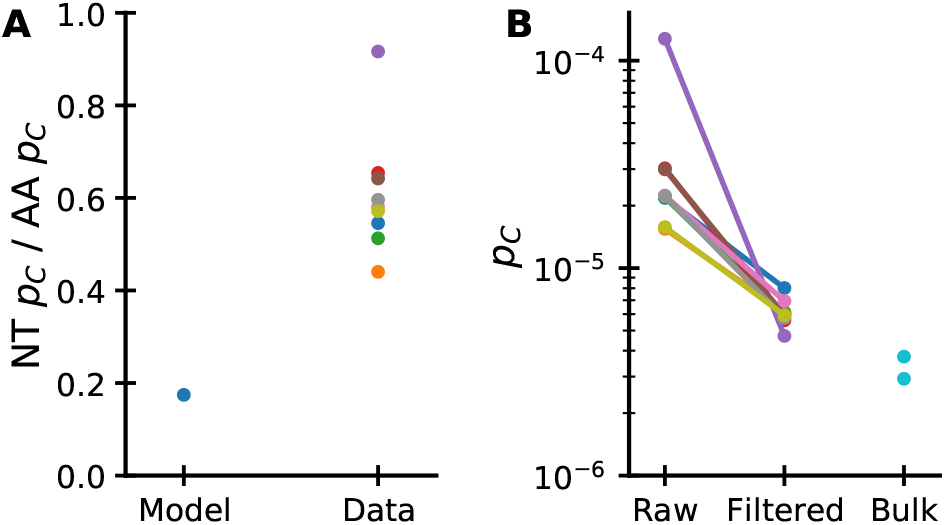
Rationale for collapsing *α* chains paired with multiple *β* chains in the paired chain data from Tanno et al. [30]. (A) The ratio between nucleotide and amino acid coincidence probabilities for *α* chains is calculated for each subject and for data generated from SONIA, a model of VDJ recombination and thymic selection. The ratio is at least two-fold higher in the data than expected suggesting substantial cross-contamination. (B) Collapsing redundant alpha chains reduces variation in amino acid coincidence probabilities across samples and makes them more comparable to those found in bulk single chain datasets (donors M and W at baseline from Minervina et al. [16]).

We next asked whether there was anything special about coincidences observed among pairs of subjects with the lowest HLA overlap. On examining the specific sequences responsible for exact coincidences in these pairs, we found that certain VJ combinations were heavily overrepresented among their *α* chains. These VJ combinations map to known signatures of noncanonical T cells with semi-invariant receptors, so called MAIT and iNKT cells [50]. These T cells have semi-invariant *α*-chains and restricted *β* chain diversity, which explains why they contribute heavily to near-coincidences. Both T cell subsets bind to non-peptide ligands, which are not presented on MHC molecules. As we are interested in identifying signatures of selection driven by pMHC binding, we exclude the VJ*α* combinations coding for these invariant T cells (TRAV1-2 paired with TRAJ12/TRAJ20/TRAJ33 for MAITs, and TRAV10 paired with TRAJ18 for iNKTs).

While these additional filtering steps have a numerically small effect on the total numbers of near-coincidences, we believe that this careful approach is warranted as some of the most interesting comparisons rest on the small numbers of exact or nearly exact coincidences.

## Appendix F: Detailed analysis of motif mixture model

In Sec. V we sketched a schematic model of TCR-pMHC binding as a string-matching problem [42, 68]. In this appendix we present a fuller account of the formulation and analysis of this model.

For simplicity we consider synthetic TCR sequences *σ* of a fixed length *k* = 6, corresponding to the number of hypervariable residues within a typical CDR3 loop. For a particular pMHC complex *p*, we assume that the binding energy depends on the residues within the TCR additively, and that each site makes a binary contribution to the energy:

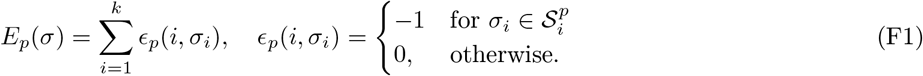

Here, for each site, 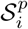 is a set of “good” amino acids that contribute to the binding of sequence *σ* to epitope *p*. A sequence is taken to bind only if *E_p_*(*σ*) = –*k*, that is to say that the amino acid at each site *i* is in the allowed set 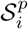. This definition of binding energy describes a random motif model, where a motif is defined as any combination of allowed amino acids at the different sites. The motif of course will vary from one epitope to another.

As in our model TCRs have fixed length we consider the simpler Hamming distance instead of edit distance, and we make two further assumptions to simplify analytical calculations: First, at each site there are an equal number *c* of allowed amino acids, 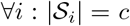. Second, background sequences are drawn from the flat distribution over all k-mers: at each site we draw independently and uniformly at random one out of the *q* = 20 amino acids. Calculating the near-coincidence histograms within the background set and within the specific set for a particular epitope then reduces to purely combinatorial exercise with the analytical results

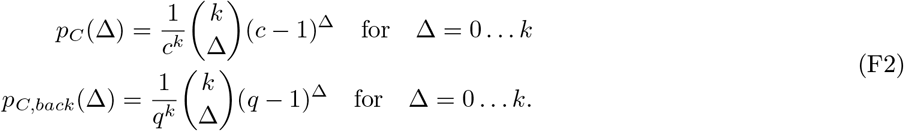

The near-coincidence enhancement factor in this model is therefore

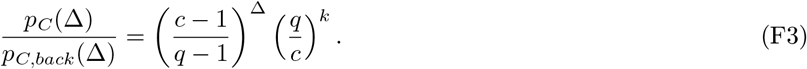

This expression reproduces the exponential falloff with Δ that is seen in real data, with the falloff rate dependent on the number of allowed amino acids *c*. To make the rate compatible with the observed factors of ten, requires that at each site there are on average *c* = 3 possible amino acids, such that the fraction of specific neighboring sequences is equal to (*c* – 1)/(*q* – 1) = 2/19 ≈ 0.1.

We next considered a mixture of motif models, where all TCRs that conform to any of *M* randomly chosen motifs are specific. Each motif defines a different binding energy function as per Eqn. F1 with independently drawn sets of allowed amino acids 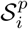. The binding energy of a TCR sequence is then taken to be the minimum over these motifs. The distribution of T cells selected by this binding energy can be approximated as a mixture of the distributions selected by the individual motifs. Applying results for coincidences in mixture distributions (derived in Appendix B), we obtain an analytical prediction for excess coincidences

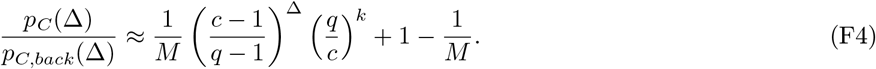

Numerical simulations of the model were performed as follows: We first draw a background set of 2 · 10^7^ TCRs of length *k* = 6. Each TCR is drawn from an independent site model, where the probability of drawing a specific amino acid is set equal to the usage frequency of amino acids found within the CDR3*α* hypervariable chains in a human blood sample from [16]. Next, we draw *M* different binding motifs, each defined by *c* = 3 amino acids drawn independently and evenly drawn from all possible amino acids. We then filter out all sequences from the background set that match the definition of any of the motifs. Finally, we calculate the coincidence probability for the specific sequences retained from the background.

